# A Multimodal Imaging Pipeline for the Discovery of Molecular Markers of Cellular Neighborhoods

**DOI:** 10.1101/2025.04.07.647675

**Authors:** Thai Pham, Melissa A. Farrow, Lukasz G. Migas, Martin Dufresne, Madeline E. Colley, Jamie L. Allen, Angela R.S. Kruse, Mark P. de Caestecker, Raf Van de Plas, Jeffrey M. Spraggins

## Abstract

Insights into tissue microenvironments are constrained by profiling cellular and molecular components independently. To address this, we developed an integrated, multimodal workflow combining imaging mass spectrometry and highly multiplexed immunofluorescence on the same sample. Custom computational pipelines enable precise spatial co-registration, facilitating direct integration of cellular and molecular profiles. CODEX-derived cellular and neighborhood masks enable mining of IMS-derived lipid data to identify molecular markers of cell types and spatial neighborhoods in human kidney tissue. We discovered distinct carnitine distributions in cortical and medullary regions, indicating metabolic heterogeneity driven by microenvironment-specific oxygen levels and energy demands. Glomerular cell-type-specific analysis found ether-linked phosphatidylcholine and sphingomyelin, species associated with slit diaphragm integrity, localized to podocytes. Immune cell neighborhoods exhibited molecular signatures consistent with cell signaling and activation pathways in damaged tissue regions. This multi-omic framework links cellular organization with molecular signatures to unveil cellular and molecular relationships underlying tissue function and pathology.

## INTRODUCTION

Cellular neighborhoods are specific regions within a tissue characterized by a consistent, repeating arrangement of different cell types that interact to influence tissue function and disease. Rather than considering individual cells in isolation, cellular neighborhoods are intermediate tissue organizational units that describe the local environment (*i.e.,* tissue microenvironment) and how cellular organization is altered during disease progression and therapeutic response. Quantifying cellular neighborhoods is essential for understanding all organ systems. For example, in the kidney, neighborhoods capture the complex architecture required for maintaining proper filtration and how specific, localized changes in cellular organization can lead to various chronic diseases^1^.

Recent advances in spatial proteomics and transcriptomics, including highly multiplexed immunofluorescence platforms such as co-detection by imaging (CODEX)^2^ and imaging mass cytometry (IMC)^3^ and transcriptomic technologies such as Xenium^4^, and Multiplexed Error-Robust Fluorescence in situ Hybridization (MERFISH)^5^, have enabled high-resolution mapping of cellular phenotypes and spatial organization. While these approaches have been transformative for defining cell identity, state, and neighborhood *in situ*, they are, in most cases, limited in their need for *a priori* target selection and probe availability. As a result, major classes of biologically important molecules, including lipids, metabolites, and post-translationally modified proteins, remain inaccessible using antibody- or probe-based spatial omics approaches.

Matrix-assisted laser desorption/ionization imaging mass spectrometry (MALDI IMS)^6–9^ enables the untargeted detection of hundreds-to-thousands of molecules directly from tissue, allowing metabolites^10^, lipids^11,12^, glycans^13^, peptides^14–16^, and proteins^17^to be mapped at cellular resolution. These molecules often lie closer to the molecular phenotype of normal and diseased tissue, reflecting metabolic state, membrane composition, signaling activity, and biochemical remodeling that are not readily inferred from protein or transcript abundance alone. By integrating MALDI IMS with targeted spatial biology technologies, a more complete systems-level view of the tissue microenvironment can be constructed, combining deep molecular profiling with cell identity, state, and spatial organization. Through these multimodal workflows, cell types, phenotypes, and neighborhood structures are defined using highly multiplexed protein or transcript imaging, while untargeted IMS reveals the underlying biomolecular landscape associated with these cellular features.

Despite this promise, multimodal IMS studies have been limited by technical incompatibilities of the combined technologies, forcing investigators to rely on lower-plexity^18,19^ or less specific microscopy techniques, or to associate data across serial tissue sections^12,18,20–22^. To date, most multimodal imaging studies integrating IMS with optical microscopy have utilized stained microscopy^23,24^, autofluorescence microscopy^12^, or low-plexity immunohistochemistry or immunofluorescence^18^ to guide segmentation and interpretation of IMS data. Such strategies have proven highly effective for mapping molecular features to larger anatomical regions^21,22^ and multicellular functional tissue units (FTUs), as demonstrated by our construction of an FTU-level lipid atlas of the human kidney^12^. These workflows established an important foundation for multimodal molecular imaging by enabling accurate and robust alignment of IMS-derived chemical information with microscopy-defined tissue features and cell types. However, the limited plexity or specificity of the microscopy technologies used in these studies did not allow for more comprehensive neighborhood information to be captured.

Recently, there have been initial efforts to integrate MALDI IMS with highly multiplexed spatial proteomics^20,25^ and spatial transcriptomics technologies^26–28^, which offer substantially greater capacity for cellular phenotyping and quantifying cellular organization. However, because of technical incompatibilities between IMS and these probe-based platforms, these approaches have relied on serial tissue sections, with data aligned indirectly across adjacent tissue slices. While serial-section approaches have yielded valuable insights, they remain fundamentally limited by section-to-section variability, tissue deformation, and a lack of spatial concordance between the multimodal measurements. As a result, molecular–cellular associations derived from serial sections are inherently probabilistic and often insufficient for resolving molecular features at the level of specific cell types, cellular neighborhoods, or single cells *in situ*.

Here, we present a fully integrated multi-omics workflow comprised of autofluorescence microscopy, high-resolution MALDI IMS, CODEX highly multiplexed immunofluorescence microscopy, and stained microscopy performed on a single tissue section. This approach retains spatial fidelity across modalities and enables a direct link between IMS-derived molecular profiles and CODEX-defined cellular phenotypes and neighborhoods. Neighborhood analysis of multiplexed immunofluorescence data revealed local cellular environments and the cell-cell interactions that drive tissue organization and function. These neighborhood definitions were subsequently leveraged to generate spatial masks to extract neighborhood-specific molecular profiles. This approach supports discovery-driven analyses that connect molecular chemistry to cellular organization and spatial context, advancing spatial biology beyond descriptive atlases toward a mechanistic, systems-level understanding of tissue function in health and disease.

## RESULTS

The optimized imaging workflow consists of autofluorescence microscopy, MALDI IMS lipid profiling, CODEX multiplexed immunofluorescence, and PAS histological staining (Fig. 1a). This workflow was performed on three human kidney samples (Methods - Table M1). MALDI IMS was collected prior to CODEX to prevent analyte delocalization, ion suppression, or loss of molecular classes. Autofluorescence images were collected before (preAF) and after (postAF) MALDI IMS imaging to facilitate the co-registration process. Following MALDI IMS and postAF acquisition, the matrix was removed and CODEX was collected. Multiplexed immunofluorescence (mxIF) following MALDI IMS has been performed previously by manual cyclic IF staining^18^. Here, that method was adapted for highly multiplexed CODEX imaging. To confirm that sensitivity was retained for staining of post-IMS samples, the signal for all 22 antibodies (Methods - Table M2) was compared to a control sample where no MALDI IMS was performed (Supplementary Fig. 1-1). Modifications to MALDI IMS typical 5 μm imaging parameters were made as highlighted in Methods - Table M3. These changes did not generate any observable alterations in the acquired data quality, as can be seen in the average spectra (Supplementary Fig. 3) and the resulting ion images (Supplementary Fig. 4-9). These changes to the instrument settings help conserve protein epitopes within the tissue section during MALDI IMS acquisition, which can range from a few hours to multi-day analyses, leading to minimal perturbation of the tissue for antibody-based imaging.

**Figure 1:**
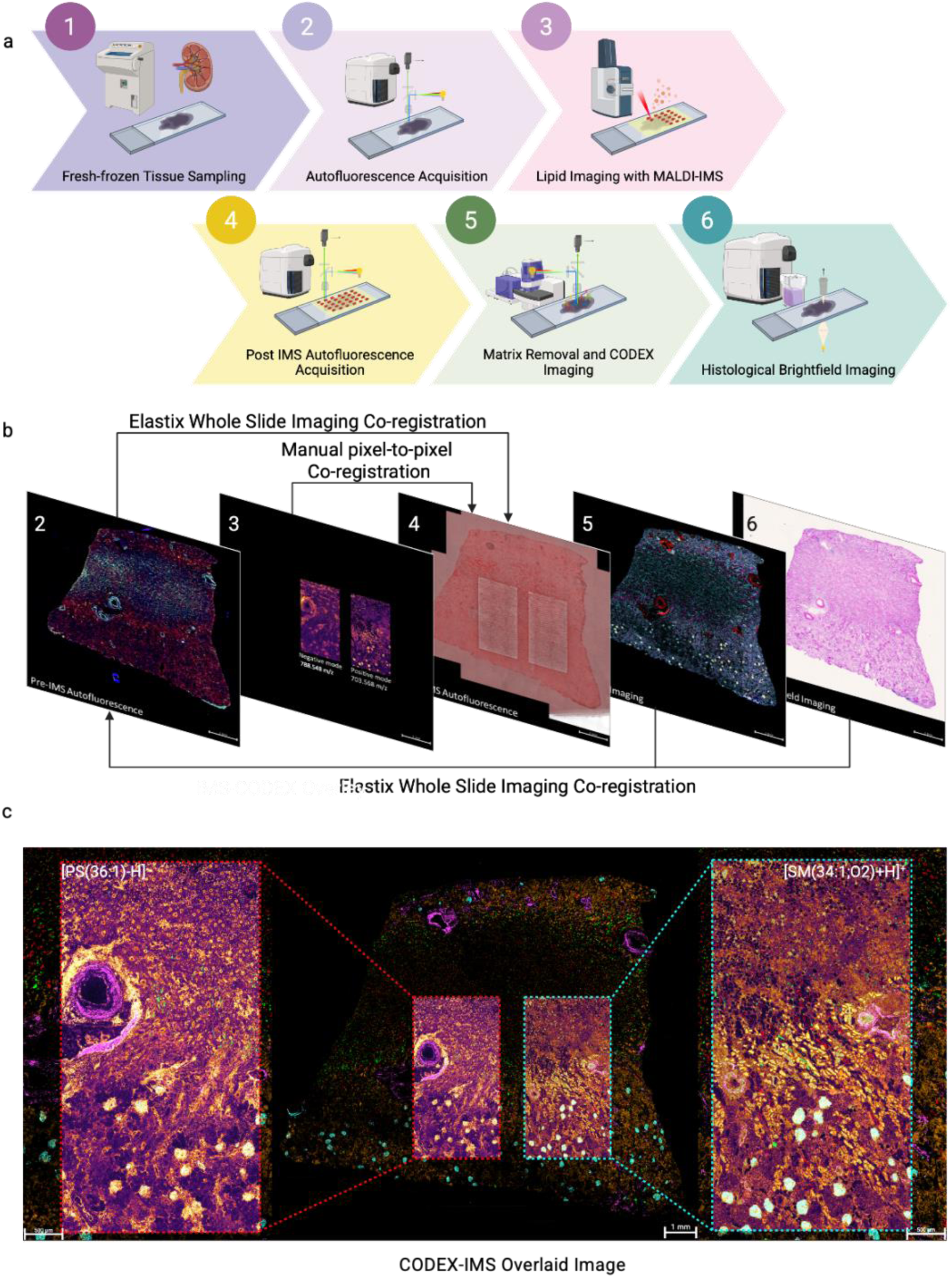
A multimodal workflow for collection and analysis of IMS-CODEX on the same tissue section. **a.** Overview of analytical pipeline for specimen processing and image acquisition. Autofluorescence was acquired on tissue sections before (pre-IMS AF) and after (post-IMS AF) MALDI IMS to support downstream image co-registration. Lipid imaging by MALDI IMS was done at 5 µm. Following matrix removal, a 22-antibody panel CODEX run was performed and PAS images were collected. **b**. Multimodal images from VAN0031 were aligned using a two-step process combining elastix and manual pixel-to pixel co-registration with the post-AF image as the central anchor point. **c**. The CODEX and IMS images were co-registered and assessed by confirming overlay of morphological features between IMS molecular distributions and CODEX cellular structures. Insets are zoomed images of [PS(36:1)-H]^−^ and [SM(34:1;O2)+H]^+^, demonstrating precision of the co-registration pipeline.

**Table M1.** Donor demographics and comorbidities.

| Sample ID | Age | Sex | Race | Height (cm) | Weight (kg) | BMI | Comorbidities |
| --- | --- | --- | --- | --- | --- | --- | --- |
| VAN0005 | 58 | F | Non-Hispanic White | 160 | 59 | 23 | Renal cell carcinoma |
| VAN0019 | 62 | M | Non-Hispanic White | 180.3 | 147.9 | 45.5 | Chronic pyelonephritis, sleep apnea, hypertension, morbidly obese |
| VAN0031 | 77 | F | Non-Hispanic White | 162.6 | 74.6 | 28.2 | Renal cell carcinoma, hypertension, type 2 diabetes |

**Table M2.** CODEX Antibody Panel and Reporters.

| Antibodies | Supplier | Catalog number | Target | Barcode | Reporter | Channel | Cycle |
| --- | --- | --- | --- | --- | --- | --- | --- |
| AQP2 | Abcam | ab230170 | Collecting Ducts (CD) | BX013 | RX013 | AF750 | 1 |
| CD7 | Abcam | ab239755 | Mature T Cell | BX025 | RX025 | AF750 | 2 |
| CD31 | Akoya | SKU 4250009 | Endothelial Cell (ECs) | BX032 | RX032 | ATTO550 | 3 |
| CD20 | Abcam | ab236434 | B Cell | BX028 | RX028 | AF750 | 3 |
| LYVE-1 | R&D System | R&D AF2089 | Lymphatic Vessel ECs | BX017 | RX017 | ATTO550 | 4 |
| CD3 | Abcam | ab245731 | T Cell | BX010 | RX010 | AF750 | 4 |
| Laminin B2 | Abcam | ab210956 | Glomerular Basement Membrane (GBM) / Vasculature Bundle/Glomerular Capsule (GNC)/Tubular Basement Membranes (TBMs) | BX041 | RX041 | ATTO550 | 5 |
| Synaptopodin | Abcam | ab282117 | Podocyte | BX006 | RX006 | Cy5 | 5 |
| Calbindin | Abcam | ab233018 | Distal Convoluted Tubule (DCT)/ Connecting | BX004 | RX004 | AF750 | 5 |
|  |  |  | Tubule (CNT)/<br>Cortical CD |  |  |  |  |
| Uromodulin | Abcam | ab223540 | Thick<br>Ascending<br>Limb of Henle<br>(TAL) | BX047 | RX047 | ATTO550 | 6 |
| CD45 | Abcam | ab269347 | Leukocytes | BX001 | RX001 | AF750 | 6 |
| Col4A1 | Abcam | ab226485 | TBM/Mesangial<br>Matrix/<br>Peritubular<br>ECM | BX014 | RX014 | ATTO550 | 7 |
| Fibronectin | Abcam | ab268022 | GNC,<br>Mesangial<br>Cells, Capillary<br>Loops,<br>Myofibroblasts | BX033 | RX033 | Cy5 | 7 |
| Ki67 | Abcam | ab279657 | Proliferation | BX043 | RX043 | AF750 | 7 |
| alpha SMA | Abcam | ab240654 | Vascular<br>Smooth Muscle<br>Cell,<br>Myofibroblasts | BX029 | RX029 | ATTO550 | 8 |
| Nestin | Abcam | ab242400 | Podocyte and<br>Endothelium | BX024 | RX024 | Cy5 | 8 |
| AQP1 | Abcam | ab178352 | Proximal<br>Tubules and<br>Descending<br>Thin Limb (DTL) | BX002 | RX002 | ATTO550 | 9 |
| Vimentin | Abcam | ab193555 | Glomerular<br>Cell /<br>Fibroblast | BX042 | RX042 | Cy5 | 9 |
| Cytokeratin 7 | Abcam | ab220804 | CD, GNC | BX005 | RX005 | ATTO550 | 10 |
| E-Cadherin | R&D System | R&D AF748 | DTL, TAL, DCT,<br>CNT and CD | BX031 | RX031 | AF750 | 10 |
| Megalin | Abcam | ab309087 | Proximal<br>Tubule | BX027 | RX027 | Cy5 | 11 |
| NaKATPase | Abcam | ab300508 | TAL, DCT, CNT,<br>CD | BX052 | RX052 | AF750 | 11 |

**Table M3.**
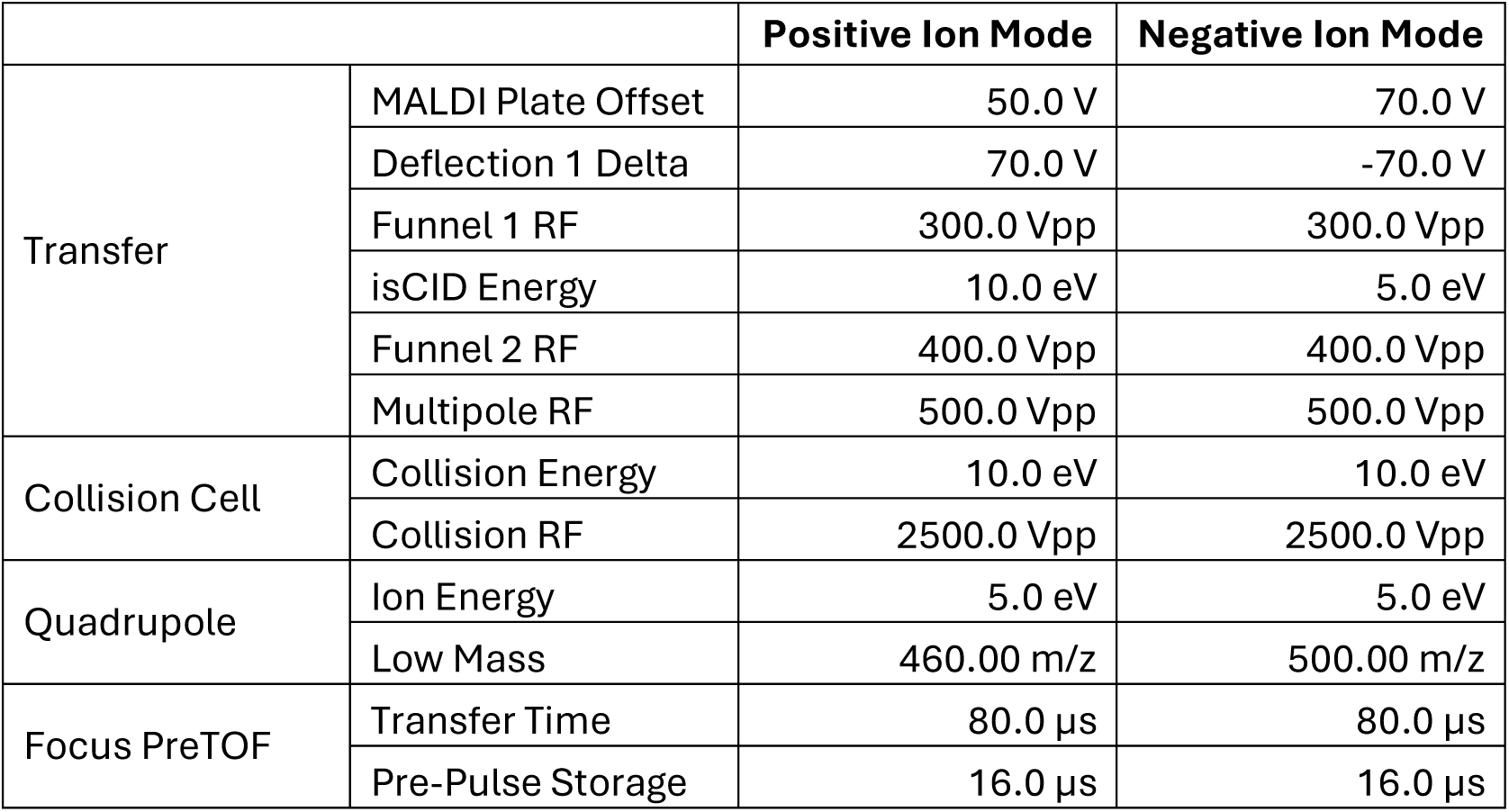
Instrument Parameter for Positive and Negative Ion Mode MALDI IMS Experiments.

|  |  | Positive Ion Mode | Negative Ion Mode |
| --- | --- | --- | --- |
| Transfer | MALDI Plate Offset | 50.0 V | 70.0 V |
|  | Deflection 1 Delta | 70.0 V | -70.0 V |
|  | Funnel 1 RF | 300.0 Vpp | 300.0 Vpp |
|  | isCID Energy | 10.0 eV | 5.0 eV |
|  | Funnel 2 RF | 400.0 Vpp | 400.0 Vpp |
|  | Multipole RF | 500.0 Vpp | 500.0 Vpp |
| Collision Cell | Collision Energy | 10.0 eV | 10.0 eV |
|  | Collision RF | 2500.0 Vpp | 2500.0 Vpp |
| Quadrupole | Ion Energy | 5.0 eV | 5.0 eV |
|  | Low Mass | 460.00 m/z | 500.00 m/z |
| Focus PreTOF | Transfer Time | 80.0 $\mu\text{s}$ | 80.0 $\mu\text{s}$ |
| | Pre-Pulse Storage | 16.0 $\mu\text{s}$ | 16.0 $\mu\text{s}$ |

The resulting multimodal datasets were co-registered using Elastix^29^, an automated framework (Fig. 1b). MALDI IMS pixels were aligned with the laser ablation marks on the postAF and registered using the image2image^30,31^ application. Then, the preAF and MALDI IMS pixel-aligned postAF were co-registered. Finally, CODEX and PAS were co-registered to the preAF using Elastix to bring all modalities into the same spatial coordinate space. Since all modalities are acquired from the same tissue section, co-registration is inherently more straightforward and capable of preserving spatial fidelity compared to aligning modalities from serial sections, where substantial perturbations such as deformations, tears, or other artifacts may be introduced. In Fig. 1c, the previously reported glomerular biomarkers PS 36:1 (*m/z* 788.544) and SM 34:1 (*m/z* 703.575)^12^ are shown overlaid with synaptopodin marker for glomeruli, confirming that the molecular species detected by MALDI IMS overlay the corresponding cellular features identified by CODEX. The co-registration of CODEX and MALDI IMS images allows for the integration of lipid analytes with cellular context. By improving the ability to link MALDI IMS derived molecular data to more granular tissue features, we can enhance discovery and improve biological interpretation of spatial lipidomics data related to specific cell types and tissue environments.

Previous lipid atlasing work in the kidney revealed lipid profiles at the functional tissue unit (FTU) level based on autofluorescence-driven segmentation^12^. Incorporating CODEX into the workflow enables spatial cell phenotyping and cell-type segmentation, which can be leveraged to mine lipid data at higher spatial resolution. Additionally, the inclusion of multiplexed immunofluorescence extends the number of FTUs detected and provides additional critical cell types, including immune cells, lymphatic vessels, and vasculature (Methods - Table M2 and Supplementary Fig. 10-13)^2,32^. Cellular and molecular patterns can be discerned from single-channel images from CODEX and MALDI IMS. Representative CODEX images for a subset of the antibody panel that identifies cell types associated with specific functional tissue units including glomeruli (synaptopodin), and proximal tubules (megalin) are shown (Fig. 2a). Similarly, ions can be detected with distributions that map to these cellular observations (Fig. 2a). SM 34:1 and SM 40:2 distributions appear to align with markers for the glomeruli and proximal tubules, consistent with previous findings^12^. We also detect carnitine species, such as CAR 20:2, that were newly found to associate with FTUs like collecting ducts (AQP2).

**Figure 2:**
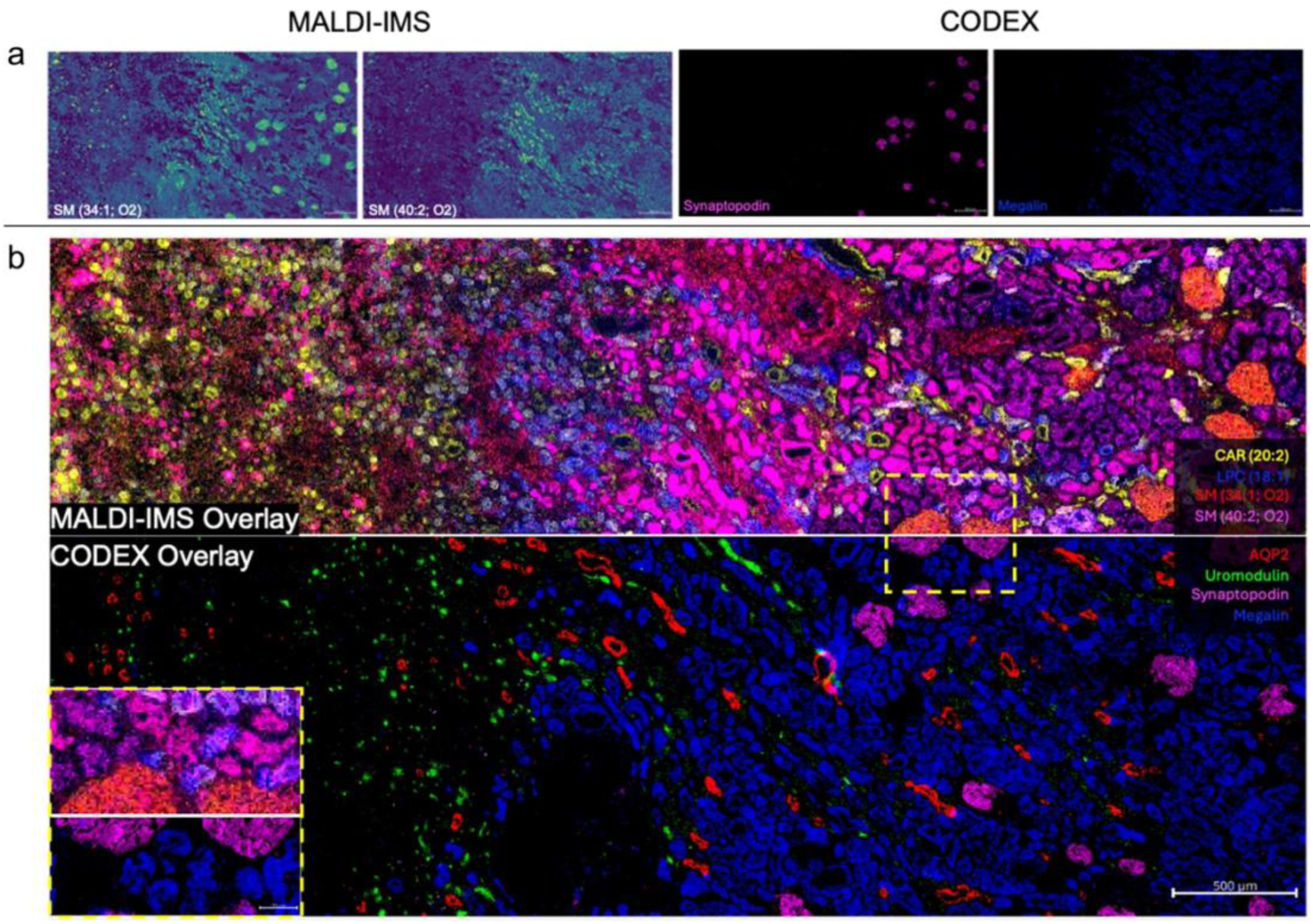
MALDI IMS and CODEX images on the same tissue section. **a.** *Selected ion images (left) illustrating spatial distributions associated with specific nephron segments, aligned with corresponding cellular marker images (right) acquired from CODEX mxIF. **b.** Overlay of MALDI IMS and CODEX images from the same tissue section, demonstrating seamless alignment between spatial metabolomic and cellular data*.

Overlaying individual channel images from CODEX provides additional detail and more in-depth cellular profiles than relying exclusively on single-marker images to identify morphological aspects of the tissue. The multiplexed nature of CODEX provides a more comprehensive view of the tissue structure and organization at the cellular level (Fig. 2b, bottom). The MALDI IMS overlay (Fig. 2b, top) shows the spatial distribution of lipids and their alignment with the tissue architecture detected by CODEX. While MALDI IMS imaging captures high-level FTU-lipid correlations, the inset image of bisected glomeruli demonstrates the increased resolution provided by the CODEX image. The ability to detect and align the lipid data from MALDI IMS with the cellular features revealed by CODEX depends on accurate co-registration across modalities from different sources and with disparate spatial resolutions, a challenge addressed by our custom computational tools that enable highly precise association of MALDI-derived molecular features with high-resolution tissue architecture.

All CODEX images (Supplementary Fig. 10-12) were processed for clustering and cell type assignment. The unsupervised machine learning approach uniform manifold approximation and projection (UMAP) produced a dimensionality-reduced representation of the data indicating 18 unique cell types can be extracted from the CODEX fluorescence intensity profiles (Fig. 3a and Supplementary Fig. 14). Cell type assignment was based on expression level profiles for each cell type (Fig. 3b). Neighborhoods were then generated (Fig. 3c) and the cell type composition of each neighborhood calculated (Fig. 3d and Supplementary Fig. 15). Ten discreet cellular neighborhoods comprised of 18 cell types were identified. Of these, FTU level neighborhoods (neighborhood A - collecting duct, B - proximal tubule, C - medullary thick ascending limb, D - cortical thick ascending limb, H - outer glomeruli, I - inner glomeruli, and J - distal tubule) were identified as well as immune (neighborhood G) and vasculature (neighborhoods E and F) specific neighborhoods. While neighborhoods share cell types, each neighborhood has a distinct population and distribution of cell types that defines it (Fig. 3d and Supplementary Fig. 15).

**Figure 3:**
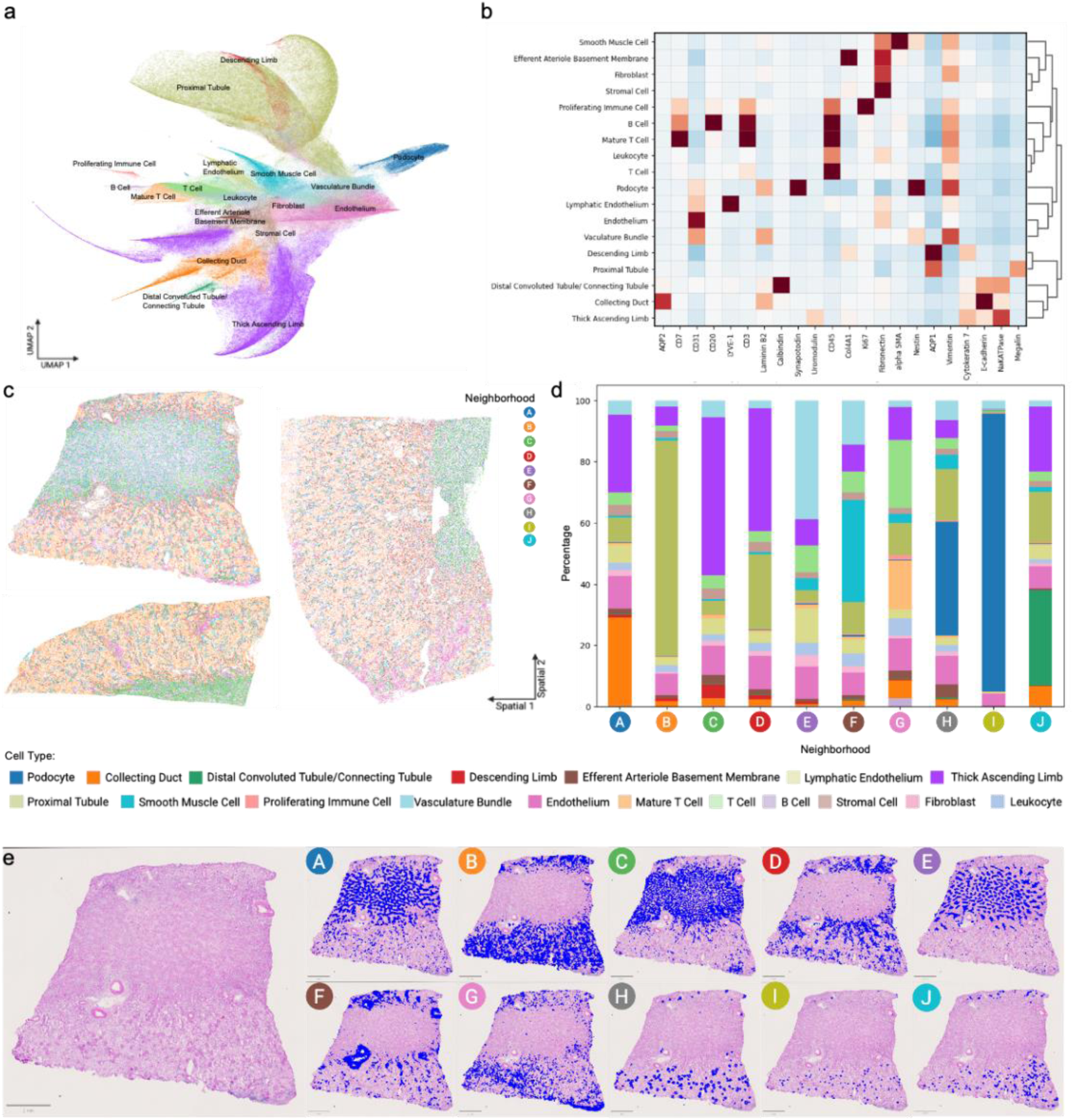
Single cell data analysis from CODEX images. **a.** UMAP visualization of integrated cell embeddings (n= 1534953), colored by cell types. **b.** Heatmap of scaled marker intensity across cell types, clustered using hierarchical dendrogram analysis. Colors represent the mean z-score of fluorescence intensity (ranging from −2 to 2). **c.** Ten cell neighborhoods from VAN0005 (bottom left), VAN0031 (top left), and VAN0019 (right), were defined based on their distinct compositions of 18 different cell types derived from protein expression data extracted from the CODEX images. **d.** The average cell type composition for each neighborhood was calculated. Each cell type is color coded as shown in key under the figure. e. Periodic acid-Schiff (PAS) stained image of human kidney tissue from donor VAN0031. CODEX-derived cellular neighborhoods are shown in a blue overlay on the PAS-stained image (Left). Each panel (A-J) represents different neighborhoods. Scale bar: 2mm.

For each sample, a neighborhood mask was generated and overlaid on the PAS stain (Fig. 3e and Supplementary Fig. 16-17). When projected onto the histological stain, each neighborhood can be seen as having a unique anatomical distribution (Fig. 3e and Supplementary Fig. 16-17). Proximal tubule (neighborhood B) and glomerular (neighborhood I) neighborhoods overlaid on donor VAN0031 highlight the differences in cell numbers and distributions of each neighborhood. While the proximal tubule and glomeruli neighborhoods are densely clustered in the cortical region, as expected, the immune neighborhood (neighborhood G) is also exclusively found in the same area. For each neighborhood, the average spectra were extracted (Fig. 4a and Supplementary Fig. 18-23). While the spectra reveal that analytes are shared across neighborhoods, the overall composition of each spectrum is unique (Fig. 4a and Supplementary Fig. 18-23), and the ion intensity of analytes varies across neighborhoods (Fig. 4b). This difference in total composition and intensity drives the specific molecular profile that defines each neighborhood. To further explore these specific molecular profiles, neighborhood masks were used to mine molecular signals by constructing classification models and interpreting them with Shapley Additive Explanations (SHAP)^33^. This framework for spatially driven molecular discovery provides a means for identifying biomarkers for each neighborhood (Fig. 4c).

**Figure 4:**
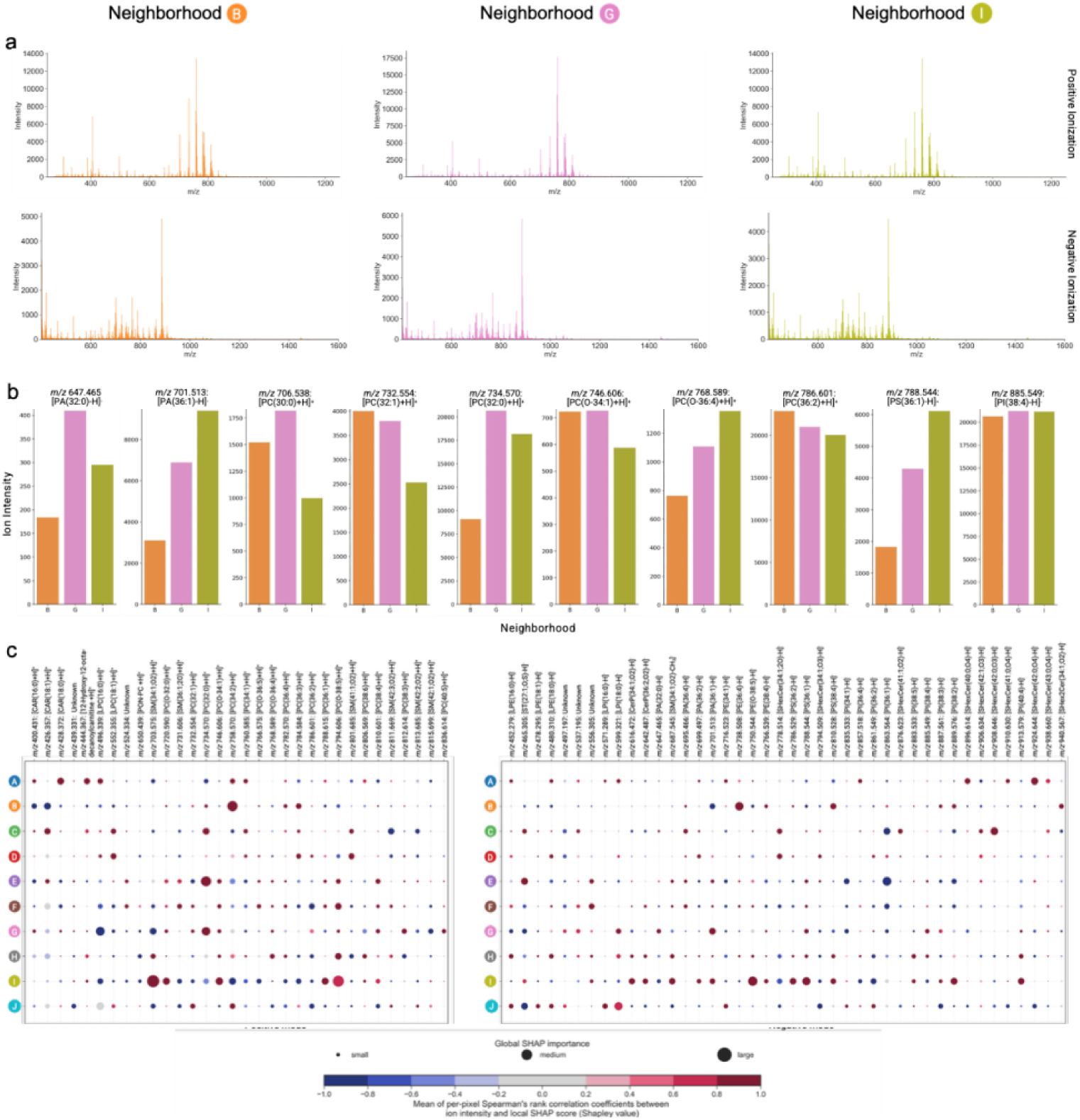
Biomolecular prolife exploration for each cellular neighborhood. **a.** Average mass spectrum of positive (top) and negative (bottom) ion mode MALDI imaging mass spectrometry data of neighborhood B, G, and I generated from donor VAN0031. **b.** Univariate ion intensities for a selection of ions that showed a strong marker profile for Neighborhood G compared to Neighborhoods B and I. **c.** SHAP bubble plot quantifying the contribution of each ion in positive (left) and negative (right) mode to the identification of the 10 neighborhoods. Each column represents an ion with ions shown in ascending m/z. Bubble size represents the magnitude of each ion’s global SHAP importance score for that neighborhood, while color indicates the direction of correlation ranging from blue (strongly anticorrelated) to red (strongly correlated).

Given that several FTU-level neighborhoods emerged, these neighborhoods were validated by comparing their molecular profiles to previously published FTU-specific data^12^. This analysis yielded results that are consistent with previous SHAP analyses. Collecting duct, neighborhood A, has exclusive correlation with SHexCer 42:0 (*m/z* 924.644), a relationship highlighted in a prior lipid atlas of the human kidney. Neighborhood B, primarily proximal tubule (PT) cells, is the only neighborhood associated with PE 36:4 (*m/z* 738.508) in line with published observations. Similarly, distal tubules were previously shown to be strongly correlated with LPE 18:0 (*m/z* 480.310), and that same observation was made for neighborhood J.

While neighborhoods dominated by FTU-specific cell types show consistency with existing datasets, they also yield new biomarkers. Neighborhood A is composed primarily of collecting ducts and the thick ascending limb (TAL), FTUs that reside in the medulla. The carnitine class is highly enriched for neighborhood A (*m/z* 400.341, *m/z* 424.341, *m/z* 426.357, *m/z* 428.372, *m/z* 442.352, *m/z* 444.367, *m/z* 448.341, *m/z* 452.372, and *m/z* 456.404) (Supplementary Fig. 26) with CAR18:0 (*m/z* 428.372) the most robust molecular marker of this neighborhood (Fig. 4d). Furthermore, when TAL is stratified into medullary (mTAL, neighborhood C) versus cortical (cTAL, neighborhood D) based on the anatomical location revealed by projection of neighborhoods onto the tissue (Fig. 3c), carnitines were found to be predominantly mTAL biomarkers. CAR 16:0 (*m/z* 400.341), CAR 18:2 (*m/z* 424.341), and CAR 18:0 (*m/z* 428.372) were strongly associated with mTAL but not cTAL (Supplementary Fig. 26). And CAR 18:1 (*m/z* 426.357) is one of the most robust mTAL biomarkers across all lipid species measured. The enrichment of carnitine species with medullary regions of the nephron could provide insight into the metabolic regulation of these specific FTUs.

Leveraging the cell type-level information provided by CODEX images, glomeruli were split into two distinct neighborhoods, with H representing outer glomeruli and I capturing inner glomerular cell types. When comparing the molecular profiles of the two glomerular neighborhoods, distinct differences were revealed. SM 34:1 (*m/z* 703.575), a known podocyte-enriched lipid^12^, is a robust biomarker of the inner glomeruli neighborhood (Fig. 4c), where podocytes predominate the cell type composition (Fig. 3d). However, its importance as a biomarker for the outer glomerular neighborhood is highly attenuated. Interestingly, there are distinct ether lipids associated with the two glomerular neighborhoods. PC O-36:4 (*m/z* 768.589) is seen in the outer glomerular neighborhood, while PC O-34:1 (*m/z* 746.606) and PC-O 32:0 (*m/z* 720.590) are more dominant in the inner glomerular neighborhood. In addition to specific ether-linked PC being associated with each glomerular neighborhood, PC 38:6 (m/z 806.569) is associated with outer glomeruli (H) while PC 36:1 (*m/z* 788.615) is a robust marker of inner glomeruli (I). Together, the outer (H) and inner (I) glomerular neighborhoods have distinct PC and ether-linked PC profiles consistent with their unique cell type populations. Additionally, PI 38:4 (*m/z* 885.549) is correlated with outer glomerular cell types (H) but anti-correlated with inner (I) cell types, suggesting distinct differences in phosphatidylinositol composition between these cell types that could impact signal transduction pathways.

In addition to FTU-level neighborhoods, vascular (E and F) and immune (G) neighborhoods were identified. All three neighborhoods (E, F, and G) are dominated by phosphatidylcholine species as positively correlated biomarkers. The vascular neighborhoods had similar PC profiles (*m/z* 746.606, *m/z* 766.575, *m/z* 768.589, *m/z* 788.615, *m/z* 794.606, *m/z* 810.601). PC O-34:1 (*m/z* 746.606), PC O-36:4 (*m/z* 768.589), PC O-38:5 (*m/z* 794.606), and PC 38:4 (*m/z* 810.601) are shared by both vascular neighborhoods (E and F) as well as with the immune neighborhood (G) while PC O-36:5 (*m/z* 766.575) and PC 36:1 (*m/z* 788.615) are only in vascular neighborhoods (E and F). Interestingly, PC 32:1 (*m/z* 732.554) was a correlative marker for neighborhoods F and G but anti-correlated with neighborhood E. The most robust biomarker for neighborhood E was also the most robust for neighborhood G, PC 32:0 (*m/z* 734.570). The other saturated PC detected in the dataset, PC 30:0 (*m/z* 706.538), shows a distinctly different association, as it is anticorrelated with neighborhood E but positively correlated with the immune neighborhood (G) (Supplementary Fig. 26). In addition, the immune neighborhood was associated exclusively with PC 38:3 (*m/z* 812.614) and PC 40:5 (*m/z* 836.614). While the enrichment of PC is a common trend across these neighborhoods, which is not seen in any of the other neighborhoods, each has its own unique PC profile that defines it.

Beyond the abundance of PC in the immune neighborhood, other lipid species are also important markers. PA 32:0 (*m/z* 647.465) is exclusively found in the immune neighborhood (G) and PA 36:1 (*m/z* 701.513) is a robust biomarker for the immune neighborhood (G). While PS 36:1 (*m/z* 788.544) is a biomarker for several neighborhoods, it is also strongly associated with the immune neighborhood (G). Phosphatidylinositol species also reveal unique associations with the immune neighborhood, as it is the only neighborhood in which both PI 38:4 (*m/z* 885.549) and PI 38:3 (*m/z* 887.561) are positively correlated markers. This specific lipid profile provides a unique molecular barcode for areas of immune cell infiltrate, potentially revealing a mechanistic link between lipids and immune cell function.

## DISCUSSION

While both MALDI IMS and CODEX provide complementary insights into tissue architecture, their markedly different spatial resolutions (5 μm pixel for IMS and 0.5 μm pixel for CODEX) prevents direct, joint analysis at the single-cell level. To overcome this limitation, we developed a workflow to acquire both modalities on a single slide, thereby maintaining spatial fidelity when co-registering the two imaging datasets. This allows us to leverage the rich cellular information from CODEX, such as cell types and spatial neighborhoods, to create masks for mining the lipid data. This approach allows us to define biologically meaningful regions to inform the automated biomarker discovery pipeline we previously developed for MALDI IMS^33^. Here, we uncovered molecular features that characterize and discriminate between biological classes, including anatomical regions, specific cell types, and spatial neighborhoods. Together, this integrative strategy links cellular organization with molecular composition, enabling the identification of biomarkers that are both spatially localized and biologically relevant across tissue regions.

The ability to pinpoint molecular signatures not only within FTUs, as we have done previously^12^, but in specific anatomical locations could be key to understanding cellular adaptations to environmental cues. Carnitines are key molecules for aerobic respiration^34^, which occurs primarily in cortical FTUs where oxygen availability is high. The enriched carnitine profile in neighborhood A, which contains CD and TAL, is intriguing, as these are FTUs typically found in the medulla, where oxygen content is highly restricted. Additionally, the stratification of TAL into medullary (neighborhood C) and cortical (neighborhood D) further highlights the increased association of carnitine species with the medullary TAL. While oxygen availability is considerably lower in this region, medullary TAL has the second highest Na/K ATPase activity across the segments of the nephron^35^. This high activity creates an immense metabolic burden on mTAL. To overcome this, the mitochondria of mTAL have adapted to the limited oxygen microenvironment and increased metabolic demands, becoming highly efficient at generating ATP under these conditions^36^. While the specific mechanism(s) that drive this adaptation have not been fully elucidated, the enrichment in carnitine availability in mTAL may be an important contributor to the increased efficiency of mitochondria observed in mTAL.

Using the CODEX-driven neighborhood analysis, the glomerular neighborhoods (H and I) were found to be enriched in ether-linked PC lipids. Interestingly, the inner and outer glomerular neighborhoods were defined by specific ether-linked PC species. For the inner glomerular neighborhood (I), which is almost exclusively made up of podocytes, two ether-linked PC species are important biomarkers, PC O-34:1 (*m/z* 746.606) and PC-O 32:0 (*m/z* 720.590). Podocytes are enriched in lipid raft regions that are crucial for maintaining the integrity of the slit diaphragm, and ether-linked lipids have been shown to be critical mediators of lipid raft organization and stability^37^. Additionally, we have previously identified SM 34:1 as a key glomerular marker that we hypothesized may also be functionally important for regulating podocyte lipid raft formation and maintenance of the slit diaphragm integrity^12^. Here, by using cell-specific markers, we see that SM 34:1 (*m/z* 703.575) is significantly more associated with the podocyte-rich inner glomerular neighborhood. It is the most robust biomarker for that neighborhood. The presence of ether-linked PC, along with SM 34:1 in the podocyte-rich inner glomerular neighborhood, points to the unique enrichment of critical lipids involved in lipid raft formation in podocytes. The ability to assign lipids to specific glomerular cell types, particularly podocytes, provides a foundation for understanding mechanisms of normal function as well as molecular drivers of disease development at the cell-type level.

Previously, studies directed at understanding lipid profiles and their role in immune cell signaling mechanisms have been restricted to cell culture models. Here, the CODEX-driven neighborhood analysis identified an immune cell neighborhood in tissues from which a molecular signature can be derived and analyzed. Similar to glomerular neighborhoods, the immune neighborhood (G) is enriched in ether-linked PC. Given that ether-linked PC are lipid raft-associated and lipid rafts are known to regulate immune cell activity, it is interesting to note that the immune neighborhood has an association with PC O-32:0 (*m/z* 720.590), PC O-34:1 (*m/z* 476.606), and PC O-36:4 (*m/z* 768.589), although their relative importance as biomarkers is diminished compared to the glomerular neighborhoods. A distinct difference between glomerular and immune neighborhoods is that the immune neighborhood is exclusively associated with all saturated PC (*m/z* 706.538, *m/z* 720.590, and *m/z* 734.570) detected in the dataset (suppl Fig. 26). Saturated PC have been shown to be specifically important in immune cell signal transduction and activity. Assembly and function of the T-cell receptor (TCR) is dependent on its recruitment to lipid rafts. A study by Zech et al showed that PC 16:0_16:0 was the most abundant PC in the TCR lipid raft domain in Jurkat T cells^38^. In addition to PC 16:0_16:0 being specifically found in the TCR domain, they also found PS 18:0_18:1 to be enriched in the TCR lipid raft region^38,39^. Both PC 32:0 (*m/z* 734.570) and PS 36:1 (*m/z* 788.544) are biomarkers for the immune neighborhood, with PC 32:0 (*m/z* 734.570) being its most robust biomarker. Lipid raft domains are pleiotropic^40^, suggesting that calibration of the lipid raft domain has to occur for fine tuning of cellular processes regulated by lipid rafts in a given cell type or cellular environment. This points to the potential for lipid rafts to be modified in a cell-type-specific manner by altering their lipid profile. Immune cells are enriched in ether-linked PC, similar to glomerular cell types, but the overall profile of lipids in the immune neighborhood is unique compared to other neighborhoods. Specifically, the identification of the immune biomarkers PC 32:0 and PS 36:1, lipids known to be critical for TCR organization and function^41^, further points to their importance as immune cell-specific regulators of signal transduction and immune cell activation.

In addition to lipids driving the organization of localized lipid raft domains for TCR assembly, they also modulate TCR activation by directly regulating enzymes involved in controlling immune cell-mediated signaling. Lipids have been shown to be critical regulators of the tyrosine phosphatase SHP1, which controls TCR activity. SHP1 recruitment to the TCR domain is mediated by binding phospholipids with high prevalence for PA species^39^. PAs were found to be strongly associated with the immune cell neighborhood (*m/z* 647.465, *m/z* 701.513), with the saturated PA 32:0 (*m/z* 647.465) being exclusively found in the immune neighborhood. Interestingly, SHP1 binds saturated PA with higher affinity^42^, consistent with our observation of PA 32:0 associated only with the immune neighborhood. Beyond the regulation of SHP1 activity, PAs are also well known, robust second messengers for signal transduction and activation in various cells. Production of PAs in immune cells is driven by the activity of phospholipase D (PLD). PLD activity is regulated by the lipid PIP_2_, which is primarily composed of PI 38:4^43^, a defining biomarker of the immune neighborhood. PIP_2_ and PA are part of a feedback loop involving PLD and PIP5K with PLD-generated PA stimulating PIP5K to produce more PIP_2_^44^. Together, these lipids drive a multitude of immune cell-specific events, including differentiation, polarization, and migration of T cells, immunological synapse organization, and cytokine production^44–46^.

This approach builds on and extends previous analysis of human kidney tissue lipid profiles linked to specific functional tissue units (FTU) of the nephron. Using CODEX-driven neighborhood analysis to mine lipid data, additional cell types and cellular interactions are detected, and unique information about cell-cell interactions, crosstalk, and niches is revealed through their cellular and molecular profiles. By leveraging the spatial fidelity provided by the multi-omic pipeline developed here, unique insight into the metabolic pathway of cortical versus medullary regions can be observed. Additionally, the enrichment of specific lipids in the immune neighborhood reveals a connection between their localization and known immune function, which has not previously been reported in tissues. More broadly, this integrated MALDI IMS–CODEX workflow represents a powerful strategy for spatial biology, enabling unbiased molecular discovery within precisely defined cellular and microenvironmental contexts. By coupling deep molecular coverage across molecular classes with high-resolution cellular phenotyping, this approach provides a powerful foundation for studying tissue organization, cell–cell interactions, and disease-associated remodeling across organ systems and biological states.

## METHODS

### Tissue Processing

Human kidney tissues were collected through the Vanderbilt Cooperative Human Tissue Network as part of cancer-related total nephrectomies as part of the Human Biomolecular Atlas Program^47,48^, following the network standard protocols and National Cancer Institute’s Best Practices for remnant surgical research material procurement. All participant provided informed consent for the collection of remnant tissue according to Institutional Review Board policies. Ethical approval for the study was granted by the Vanderbilt University’s IRB (IRB #210190), which oversees human subject involved research to ensure compliance with ethical standard. Details of donor characteristics, including age, sex, race, body measurements, and health conditions, are provided in **Table M1**.

Human tissues were flash frozen and stored at −80 °C. Tissues were sectioned at 10 µm thickness using a CM3050S cryostat from Leica Microsystems (GmbH, Wetzlar, Germany) and thaw mounted on ITO-coated microscope glass slides. The tissue sections were then washed with 4 solutions of isotonic ammonium formate (150 mM) for 45 in each solution and dried in a vacuum desiccator for 20 minutes. Samples were then vacuum sealed and stored in a −80°C freezer until analyzed. Autofluorescence images of each section were captured using standard DAPI, eGFP, and DSRed filters on a Zeiss AxioScan.Z1 slide scanner (Carl Zeiss Microscopy GmbH, Oberkochen, Germany) with a Colibri7 LED light source.

### CODEX Multiplexed Immunofluorescence and PAS-Stained Microcopy

Following the acquisition of post-IMS autofluorescence images, tissue slides were submerged in HPLC-grade acetone for 2 minutes, repeated five times, to thoroughly remove any residual chemical matrix^49^. The slides were then air-dried for 2 minutes before proceeding with the standard staining protocol detailed by Akoya Biosciences.

Briefly, the staining process began by hydrating the sample using the Akoya Hydration Buffer, followed by fixation in 16% methanol-free formaldehyde (Pierce™), diluted at a 1:9 ratio (v/v) with Akoya Hydration Buffer, for 10 minutes. This was followed by a 20-minute buffer exchange using Akoya Staining Buffer prior to the addition of conjugated antibodies. The slides were then incubated with the antibodies for 3 hours. Details of the antibodies used are provided in **Table M3**. After incubation, the slides were washed twice by submersion in Akoya Staining Buffer for 2 minutes each, followed by a 10-minute fixation in 16% methanol-free formaldehyde (Pierce™), diluted 1:9 (v/v) with Akoya Storage Buffer. This was followed by three washes with 1X PBS. To immobilize the DNA barcodes conjugated to the antibodies, the slides were submerged in ice-cold HPLC-grade methanol for 5 minutes, followed by three additional washes with 1X PBS. A final fixation step was performed using 20 μL of Akoya Fixative Reagent diluted in 980 μL of 1X PBS, followed by three more PBS washes. Finally, flow cells were assembled, and the slides were imaged using the Akoya Phenocyler and Akoya Fusion 2.0 system.

The reporter plate for this experiment was prepared according to the standard protocol provided by Akoya Bioscience. Reporter details were described in **Table M3**. All Akoya buffers used in this protocol are included in the Akoya Sampling Kit (Akoya SKU: 7000017). Once CODEX images are obtained, tissue slides are submerged into HistoChoice Clearing Agent for 24 hours to remove the flowcell before performing Brightfield PAS stain^50^.

### MALDI Matrix Sublimation

Samples were prepared using an in-house designed sublimation apparatus as described previously^51^ and similar to the HTX sublimate platform (HTX Technologies, Chapel Hill, NC). In short, the prototype sublimation apparatus is filled with 2 mL of a 3 mg/mL solution of matrix^52^. Sublimation is then performed at 200°C for 15 min at <200 mTorr, completely transferring all of the matrix onto the sample while keeping the slide at −78°C using dry ice slurry. The final matrix density on the sample slide was 0.25 µg/mm^2^. Matrix surface concentrations were determined by weighing the tissue-mounted ITO slides before and after deposition of the chemical matrix. The matrix coated slide then underwent an additional step of thermal annealing step where each slide is placed on a hot plate for 30s at 100°C before analysis to improve crystallization of the chemical matrix layer^52^.

### MALDI Imaging Mass Spectrometry

Matrix-assisted laser desorption/ionization (MALDI) imaging mass spectrometry (IMS) was performed on a Bruker MALDI timsTOF fleX mass spectrometer^53^(Bruker Daltonics, Billerica, MA) equipped with a microgrid system in positive and negative ion mode. aser power was set at 3.2% of the maximum output of the frequency tripled Nd;YAG laser (355 nm) using 20 shots per pixel. Data acquisition was performed using timsControl 5.1 and flexImaging 7.4 (Bruker Daltonics, Billerica, MA). The image pixel size (*i.e.,* pitch) was set to 5 µm, and laser spot size was confirmed to be less than 5 µm with brightfield microscopy. Instrument-specific parameters are available in **Table M2.** MALDI IMS visualization was performed using SCiLS lab 2023c Pro. After acquiring the MALDI IMS data, autofluorescence and monochromatic brightfield images of the same tissue sections were collected before removing the matrix using a Zeiss AxioScan.Z1 fluorescence slide scanner with the previously mentioned eGFP and DSRed filters. These images were used to enable high accuracy image co-registration between IMS and the microscopy modalities (See Multimodal Image Registration below)^54^.

### Multimodal Image Registration

MALDI IMS and microscopy images were registered in a two-step process. Microscopy images were co-registered using the *elastix* framework integrated into the in-house developed image2image (v0.1.7)^30,31^ application. We use *rigid* and *affine* transformations for this step, as substantial differences (*e.g.,* deformations, tears, or other artifacts) are not expected within the same tissue section (all microscopy modalities and IMS were collected on the same tissue section). For example, for each tissue section, we co-register the PAS to post-IMS AF, the CODEX to post-IMS AF via PAS, and pre-IMS AF to post-IMS AF via CODEX. The indirect registration workflow permits accurate registration of imaging modalities that have different characteristics (*e.g.,* pixel sizes). All registered whole-slide images are stored in the vendor-neutral pyramidal OME-TIFF format and retain their original spatial resolution. The MALDI IMS pixels were aligned to laser ablation marks as measured by the post-IMS AF image using image2image (v0.1.7) application. An affine transformation is computed between the post-IMS AF and IMS pixels. The transformation can then be applied to auxiliary data obtained from *e.g.,* pre-IMS AF (FTU segmentations) or CODEX (cell-type assignments or neighborhood segmentations), and the microscopy-level information can be transferred to IMS coordinate space. Similarly, the inverse affine transformation can be used to display IMS images on top of co-registered microscopy modalities.

### CODEX Single Cell Segmentation

Single-cell segmentation was performed using the Cellpose plugin (version 2.2.2)^55^ integrated into QuPath (version 0.5.0)^56^. Segmentation was based on a nuclei channel stained with the nuclear marker 4’,6-diamidino-2-phenylindole (DAPI). The Cellpose2D algorithm was implemented using the cyto3 pretrained model with the following settings: pixel size of 0.5 µm for detection resolution, tile size of 1024 pixels for efficient processing, and a cell expansion of 5.0 µm to approximate cell boundaries from nuclear masks. Intensity measurements were calculated across all cellular compartments. The script was executed on selected annotations within the QuPath project, with error handling for cases lacking a parent object. Post-processing steps, such as filtering small objects, were deferred to separate scripts for validation.

### CODEX Cell Type Clustering

Multiplexed fluorescence intensity measurement data extracted from individual segmented cells were analyzed using SCANPY (version 1.10.0)^57^. For each sample, data were normalized, and a batch number was assigned corresponding to the sample ID. Principal component analysis (PCA) was performed on individual sample datasets. Data from 1.2 million cells across three samples was retrieved by integrating 3 samples together using the BBKNN (batch-balanced k-nearest neighbors) algorithm within SCANPY to correct for batch effects. A neighborhood graph was constructed, and Uniform Manifold Approximation and Projection (UMAP) was calculated to visualize cell clusters in two-dimensional space. Cells were clustered using the Leiden algorithm, and cell types were defined based on the fluorescence intensity profiles of specific cell markers.

### CODEX Neighborhood Analysis

Cellular neighborhoods were defined by identifying the 30 nearest neighboring cells for each individual cell. The frequency of each cell type within these neighbor windows was recorded. Cells with similar neighbor window profiles were grouped into neighborhoods, reflecting spatial proximity. This analysis paralleled the cell type clustering workflow, utilizing SCANPY’s neighborhood graph and clustering tools. Each neighborhood mask was generated and exported as a separate .geojson file for downstream mass spectrometry analysis.

### MALDI IMS Data Pre-Processing

MALDI IMS data were exported from the Bruker timsTOF file format (.d) to a custom binary format. Each pixel/frame contained centroid peaks spanning the entire acquisition range. These were reconstructed into a pseudo-profile mass spectrum using Buker’s SDK (v2.21)^58^. The data were *m/z*-aligned using at least six peaks commonly present in the majority of pixels, utilizing the msalign^59,60^ library (v0.2.0). The mass axis of each dataset was calibrated using a minimum of four theoretical masses to achieve an accuracy of approximately ±1 ppm. Subsequently, the MALDI IMS data were normalized using a total ion current (TIC) approach, and an average mass spectrum was computed based on all pixels. This average spectrum underwent peak-picking and deisotoping, resulting in 376 and 413 peaks in negative and positive ion modes, respectively. Each peak list was then used to extract ion centroid data and was integrated within a window of approximately ±5 ppm. This value varies slightly depending on the bin spacing in the specific mass spectrum.

To enable neighbourhood analysis, the CODEX-based masks were transformed to align with the IMS coordinate system using the previously mentioned inverse affine transformation.

### Supervised Machine Learning and Shapley Additive Explanations

Neighbourhood annotations obtained from the CODEX experiments were used as classes to build eXtreme Gradient Boosting (XGBoost) classification models designed to recognize one of the available classes. We take the one-versus-all approach to multi-class classification, where each classification task aims to differentiate the positive class against all other pixels in the dataset. To identify which molecular species (*i.e.,* detected ion by the mass spectrometer) have a marker-like relationship to each mask, we employed Shapley Additive Explanations (SHAP). SHAP quantifies the importance of each ion in recognizing the mask within the classification model, providing both experiment-wide (global) and per-pixel (local) importance scores.

Global SHAP importance scores rank all molecular species by decreasing relevance for recognizing a specific class, highlighting a set of highly discriminative molecules that represent potential biomarker candidates. Conversely, local SHAP importance scores assess the direction of relevance (positive or negative monotonic correlation) and evaluate the significance of the relationship between an ion species’ intensity and a pixel’s likelihood of belonging to a class.

The classification models were trained using a 67%/33% train/test split and implemented with the scikit-learn^61^ (v1.3.0), XGBoost^62^ (v2.1.1), and SHAP^63^ (v0.46.0) libraries in Python (v3.9.18). Given that some masks cover less than 5% of the dataset, leading to extremely unbalanced data, we used imbalanced-learn^64^ (v0.11.0) to adjust the dataset balance by over-sampling the minority class with synthetic data augmentation and under-sampling the majority class through random subsampling. In order to ensure the reproducibility of the biomarker workflow, we train 10 independent XGBoost models for each classification task. Each XGBoost model is initialized using a different random seed, and the models are trained on slightly different training dataset (since different random seed is also used during the train/test split and data balancing).

We ranked the SHAP scores in descending order of importance, creating a shortlist of molecular species that may serve as useful biomarker candidates. In the summary bubble plots, the size of each marker represents the global SHAP importance score for each ion (column) for a given classification task (row). Additionally, we determined whether a biomarker candidate is positively or negatively correlated with a given ion species by calculating the Spearman rank-order correlation coefficient (*p*) between the mean-centred intensity and the Shapley values of each molecular ion. The Spearman correlation coefficient ranges from −1 to 1, where magnitudes greater than 0.2 are considered significant. In the bar charts and summary bubble plots, the marker color corresponds to this correlation coefficient for each ion species, illustrating the strength and direction of their relationships.

Due to the multi-sample nature of the data, the image centroids were first normalized using TIC and subsequently inter-normalized using the median change approach described by Veselkov *et al*^65^. and implemented in *ims-utils* library (v0.1.6)^66^.

### LC-MS/MS Methods and Molecular Annotation

To create a custom, human kidney lipid database for annotating MALDI IMS data, lipid extracts were analyzed from 84 human kidney tissue donors, including the three donor tissues used in these IMS-CODEX studies. All tissues were cryosectioned at 10 µm thickness into Thermo glass shell vials. Three sections were collected per patient. 5 µL of 100 µg/mL Avanti Equisplash, Avanti standard C24:1 mono-sulfo-galactosyl(B) ceramide-d7 (d18:1/24:1), and Avanti 18:2 Cardiolipin-d5 Lipids were added as internal standards. Lipids were extracted with MTBE^67,68^ and the top layer was dried to completion then resuspended in 400 µL of methanol. A pooled QC was made by collecting 5 µL from each patient. LC-MS/MS was performed with a Waters Acquity Premier UHPLC (Waters, Milford, MA) interfaced with a Bruker timsTOF Pro2 mass spectrometer (Bruker Daltonics, Billerica, MA). Reversed-phase LC separations occurred on a 2.1 mm x 100 mm Waters Premier CSH-C18 column following the gradient profile outlined in **Table M4**. Mobile phase A consisted of 60:40 ACN:H2O with 10 mM ammonium formate and 0.1% formic acid. Mobile Phase B consisted of 90:10 IPA:ACN with 10 mM ammonium formate and 0.1% formic acid. The timsTOF Pro2 mass spectrometer utilized parallel accumulation-serial fragmentation (PASEF) and MS/MS stepping to obtain the collision cross section (CCS) of each species to permit the simultaneous measurement of both low *m/z* fragments and higher molecular weight intact ‘parent’ lipids. LC-MS/MS instrument-specific parameters can be found in **Table M5**. Analysis was performed with MS-DIAL version 4.90 and all lipid annotations were blank filtered, retention scored, and referenced matched with the MS-DIAL combined database. Lipid annotations for individual patient samples analyzed in this study were obtained from the specific patient analyses against the pooled QC for 84 patients. Lipids annotated by LC-MS/MS were removed of adducts (e.g. HCOO-) prior to mass matching by accurate mass for IMS data. MALDI IMS fragment ions and gas-phase adducts were determined by on-tissue fragmentation matching to LC-MS/MS fragmentation patterns.

**Table M4.** Elution profile for lipids on a Waters Premier UHPLC.

| Time (min) | %A |
| --- | --- |
| Initial | 100 |
| 1.00 | 80 |
| 5.00 | 40 |
| 17.00 | 1 |
| 19.00 | 1 |
| 20.00 | 1 |
| 21.00 | 100 |
| 25.00 | 100 |

**Table M5.** Instrument parameters for LC-MS/MS in positive and negative ion modes on the timsTOF Pro2 mass spectrometer with 4D-PASEF.

|  | Negative Ion Mode |  | Positive Ion Mode |
| --- | --- | --- | --- |
| Mass Range (m/z) | 50-2000 |  | 100-1350 |
| Capillary | 3800 V |  | 4500 V |
| Dry Gas | 8 l/min |  | 10.0 l/min |
| Dry Temp | 220 °C |  | 220 °C |
| Nebulizer | 2.0 bar |  | 2.2 bar |
| 1/K<sub>0</sub> Start | 0.55 |  | 0.55 |
| 1/K<sub>0</sub> End | 2.20 |  | 1.85 |
| MS/MS Stepping | Scan 1 | Scan 2 | No Stepping |
| Collision RF | 450 V | 2200 V |  |
| Transfer Time | 25 µs | 120 µs |  |
| Pre-Pulse Storage | 5 µs | 12 µs |  |
| CE (eV) @ 1/K<sub>0</sub> Start | 40.0 | 40 | 30.0 |
| CE (eV) @ 1/K<sub>0</sub> End | 40.0 | 55.0 | 30.0 |

## ACKNOWLEDGEMENTS

We thank the Hubmap data curation and ingest team, particularly B. Honick, P. Blood, and J. Silverstein, for the support.

## Author contributions

Conceptualization: T.P., M.A.F., A.R.S.K., R.V.d.P. and J.M.S. Investigation: T.P., M.A.F., L.G.M., M.D., M.E.C., J.L.A., A.R.S.K., M.P.d.C., R.V.d.P. and J.M.S. Formal analysis: T.P., M.A.F., L.G.M., M.D., M.E.C., M.P.d.C., R.V.d.P. and J.M.S. Funding acquisition: M.A.F., R.V.d.P., and J.M.S. Project administration: M.A.F., R.V.d.P., and J.M.S. Supervision: M.A.F., R.V.d.P., and J.M.S. Visualization: T.P., M.A.F., L.G.M., J.M.S. Data curation: T.P., M.A.F., L.G.M. Validation: T.P., M.A.F., M.E.C., M.P.d.C., R.V.d.P. and J.M.S. Software: T.P., L.G.M., R.V.d.P. Writing—original draft: T.P., M.A.F., J.M.S. Writing—review and editing: T.P., M.A.F., L.G.M., M.D., M.E.C., J.L.A., A.R.S.K., M.P.d.C., R.V.d.P. and J.M.S.

## Data and materials availability

All data needed to evaluate the conclusions in the paper are present in the paper and/or the Supplementary Materials.

