## Supplemental Figures for "A Multimodal Imaging Pipeline for the Discovery of Molecular Markers of Cellular Neighborhoods"

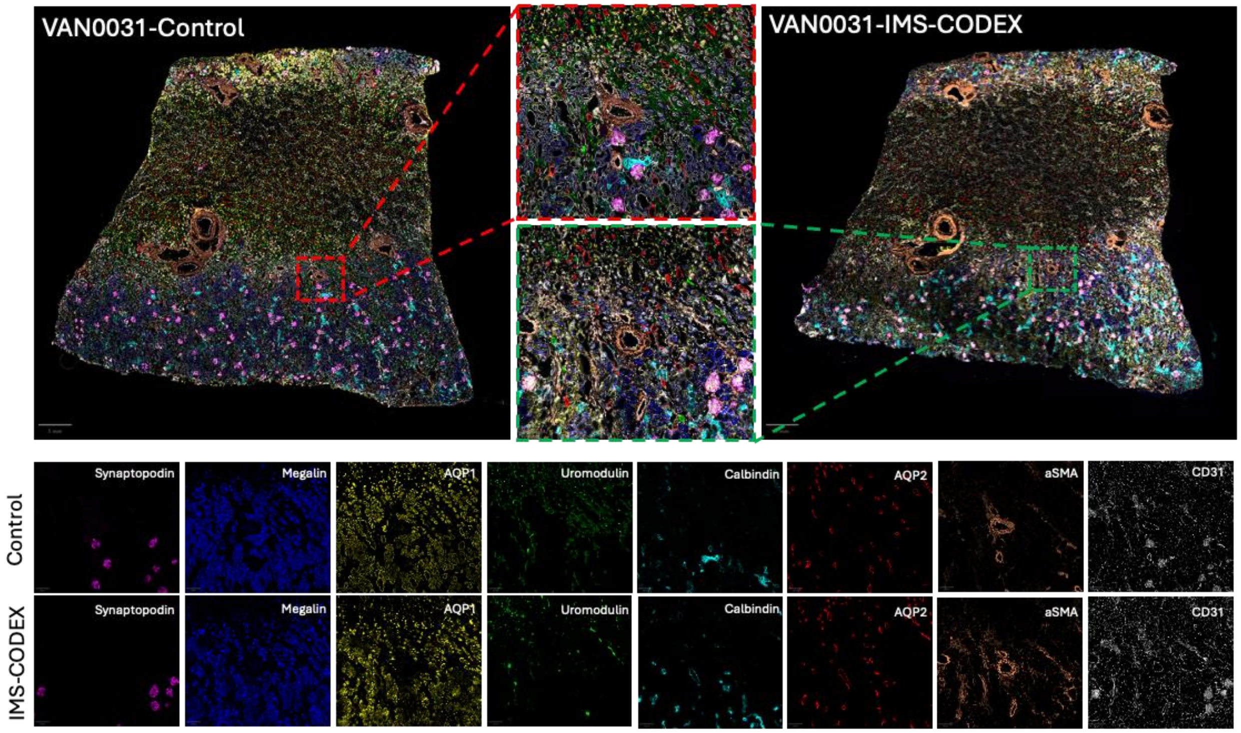


***Supplementary Figure 1:*** *MS–CODEX (right) conditions show consistent tissue morphology and marker localization across modalities. Magnified insets highlight well-preserved alignment and marker distribution in both datasets. The color of each single-channel image in the bottom panel corresponds to the same marker color shown in the full-tissue composite images above. Single-marker panels (Synaptopodin, Megalin, AQP1, Uromodulin, Calbindin, AQP2, αSMA, and CD31) demonstrate comparable signal intensity, spatial resolution, and structural clarity between the control and IMS–CODEX images.*

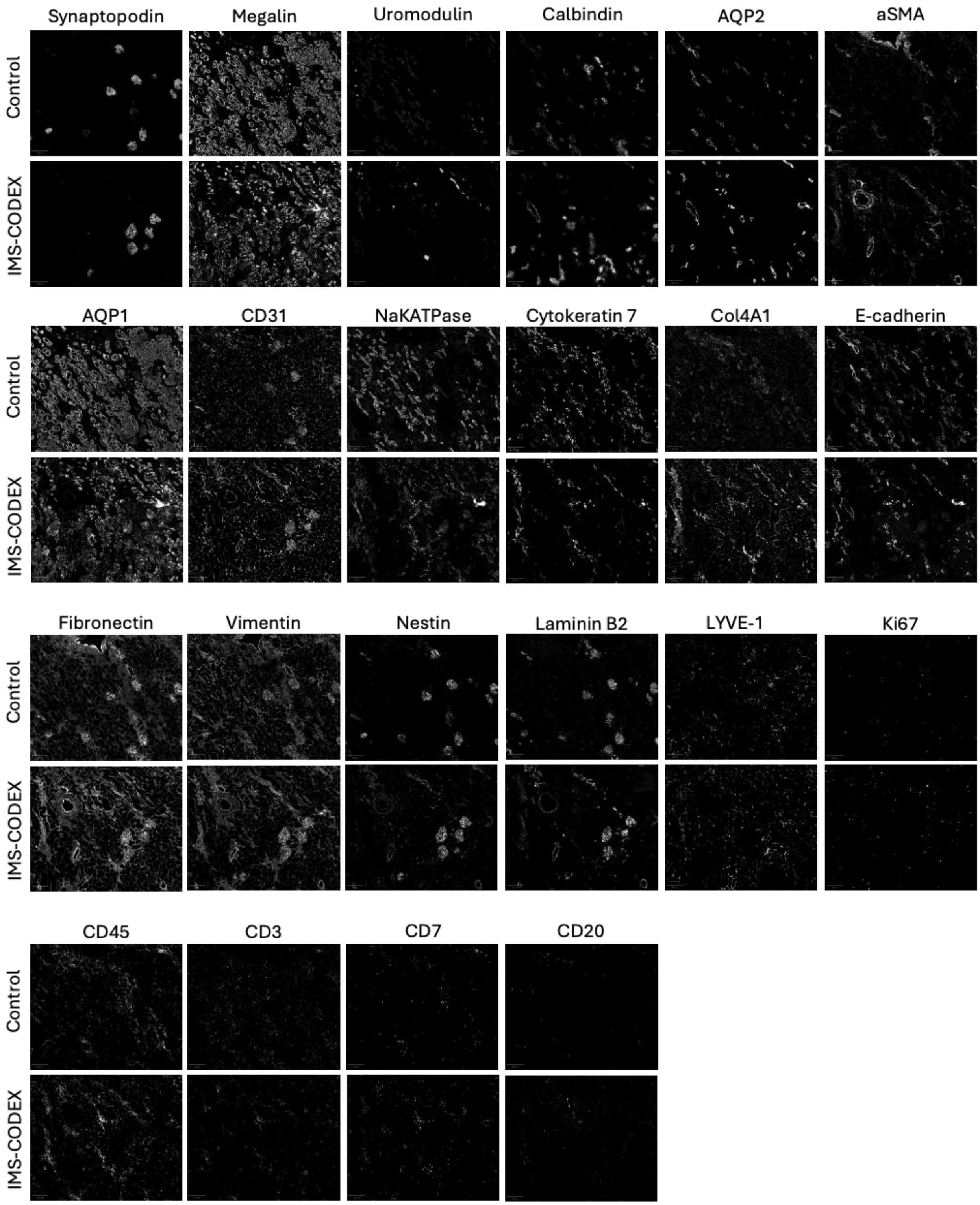


***Supplementary Figure 2:*** *Comparison of 22-marker antibody signals between control and IMS–CODEX conditions. Magnified grayscale images from kidney tissue (sample VAN0031) show all 22 antibody channels used in the imaging panel under control and IMS–CODEX conditions.*

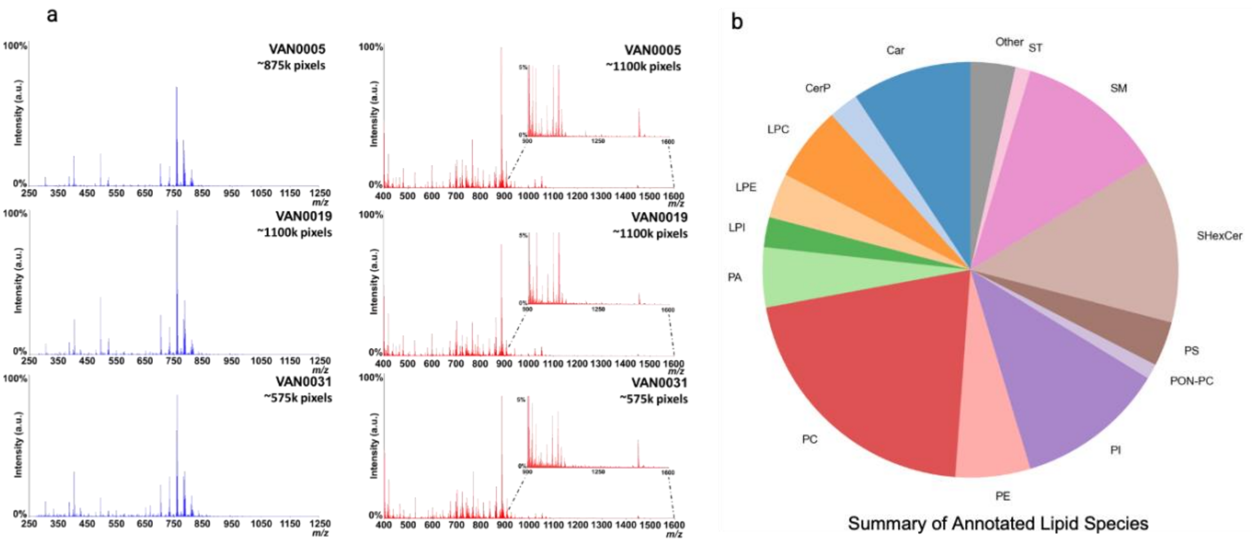


***Supplementary Figure 3: a.*** *Average mass spectrum of positive (left-blue) and negative (right-red) ion mode MALDI imaging mass spectrometry data generated from VAN0005, VAN0019, and VAN0031.* ***b.*** *Pie chart illustrating the distribution of annotated lipid species identified in the study.*

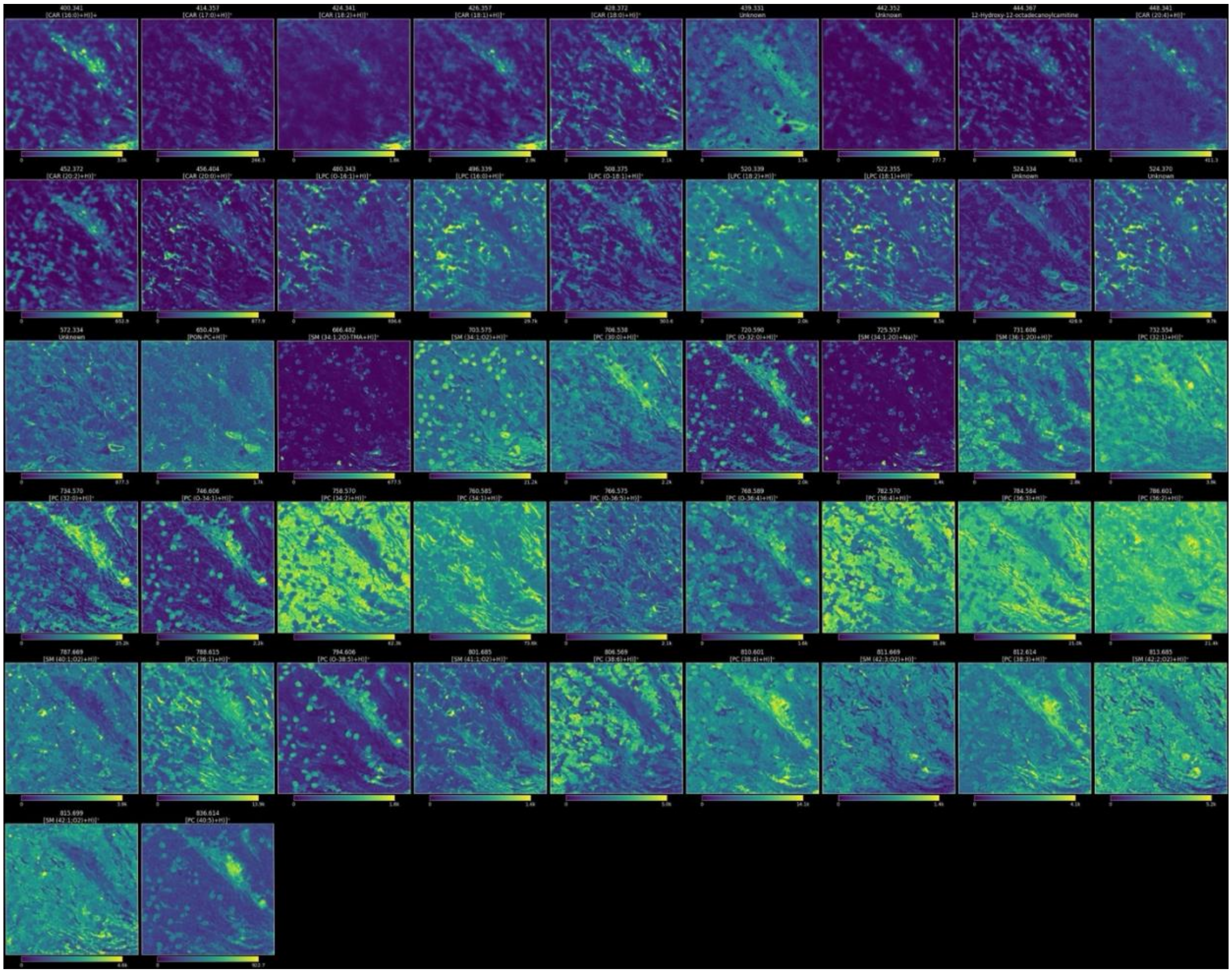

***Supplementary Figure 4:*** *Mosaic of positive ion mode MALDI imaging mass spectrometry data generated from donor VAN0005.*

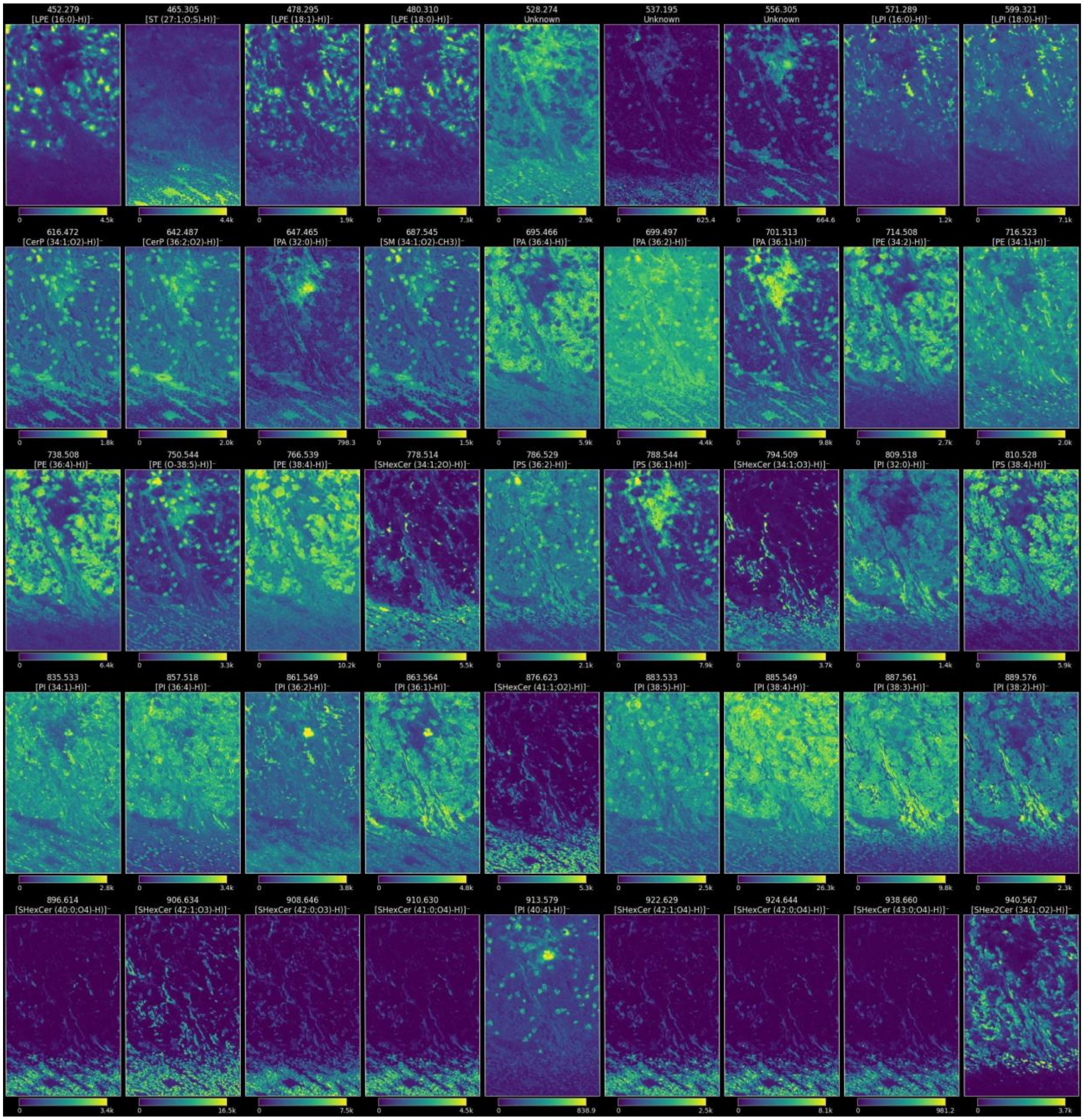


***Supplementary Figure 5:*** *Mosaic of negative ion mode MALDI imaging mass spectrometry data generated from donor VAN0005.*

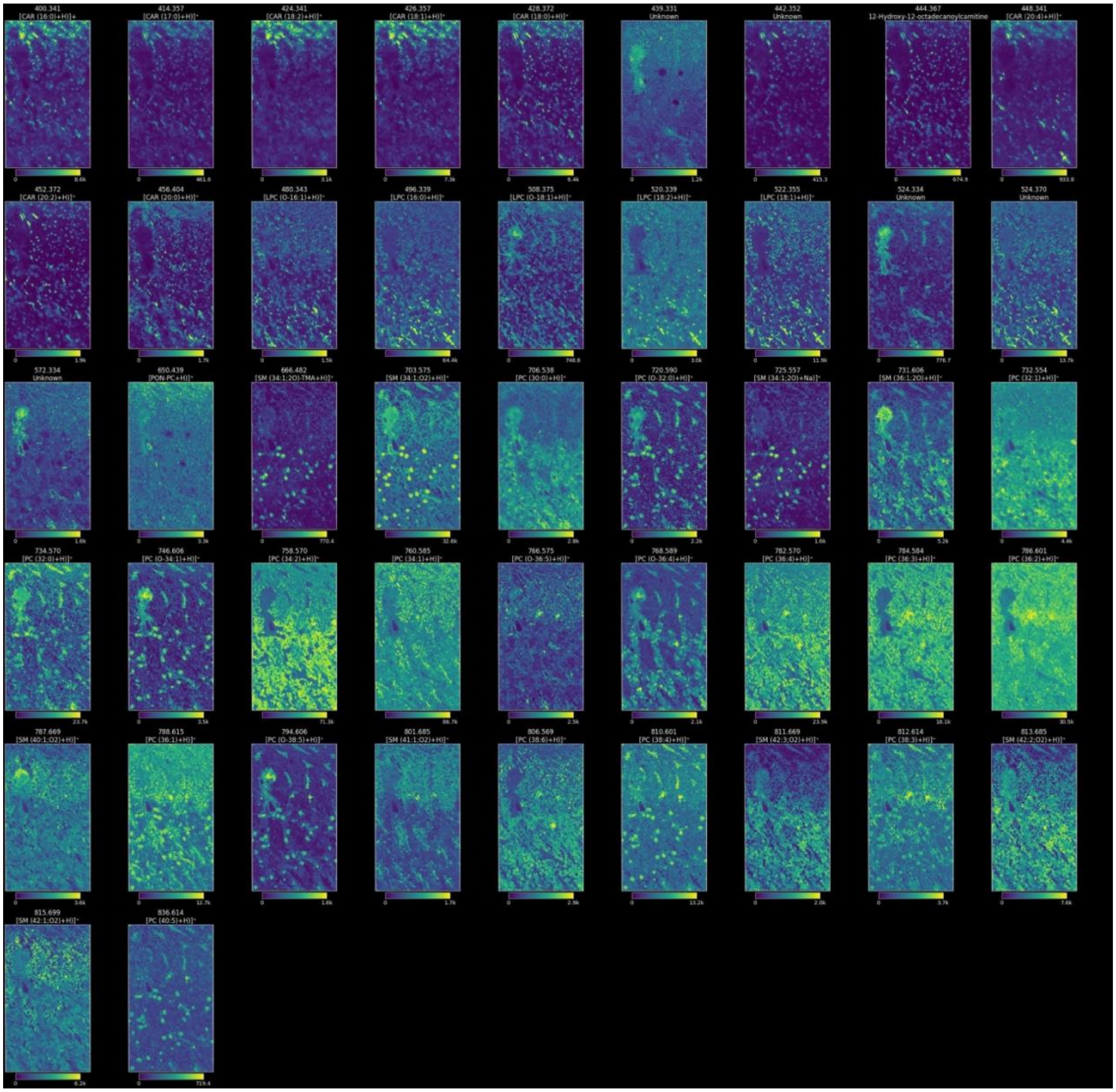


***Supplementary Figure6:*** *Mosaic of positive ion mode MALDI imaging mass spectrometry data generated from donor VAN0019.*

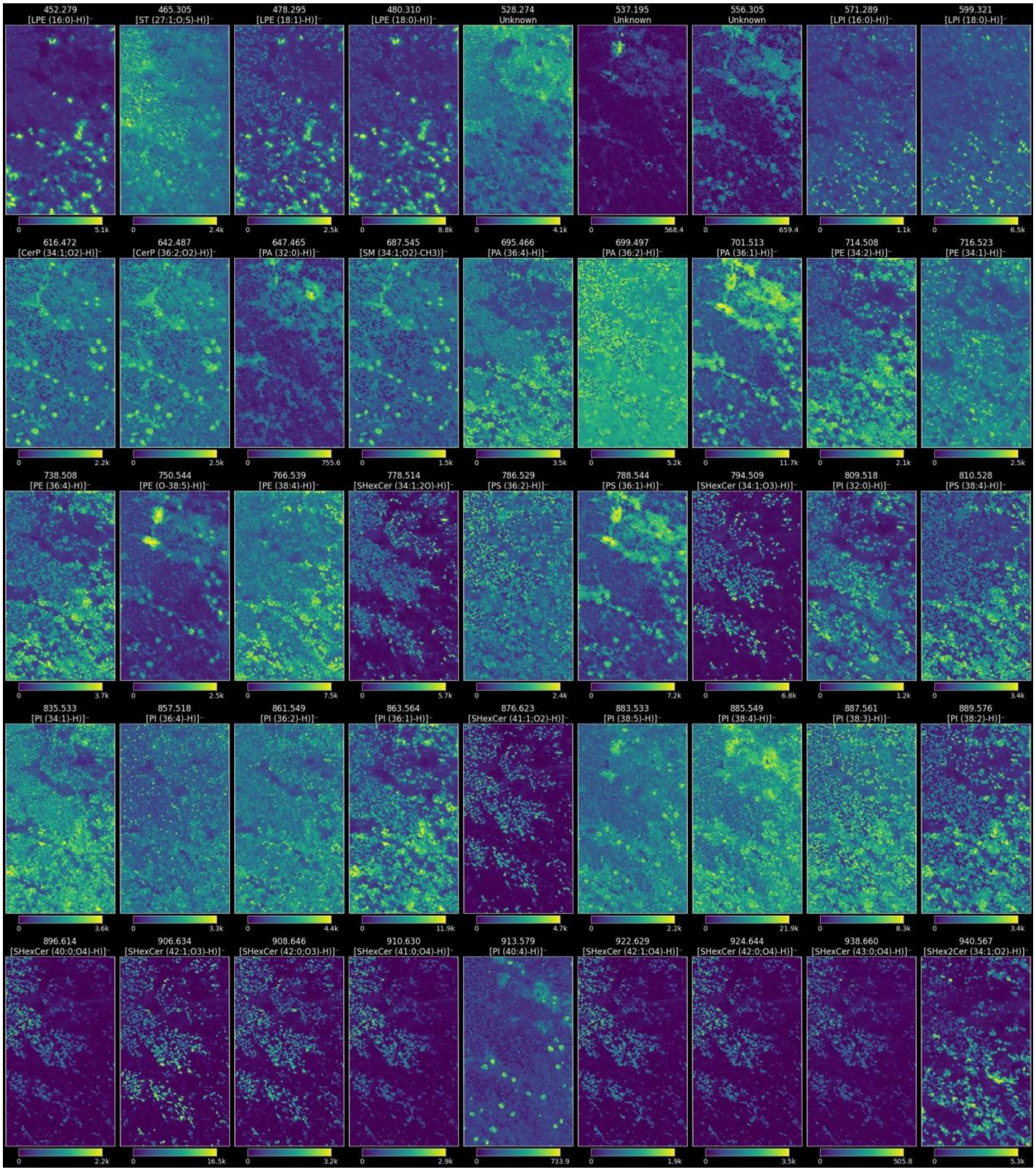


***Supplementary Figure 7:*** *Mosaic of negative ion mode MALDI imaging mass spectrometry data generated from donor VAN00019.*

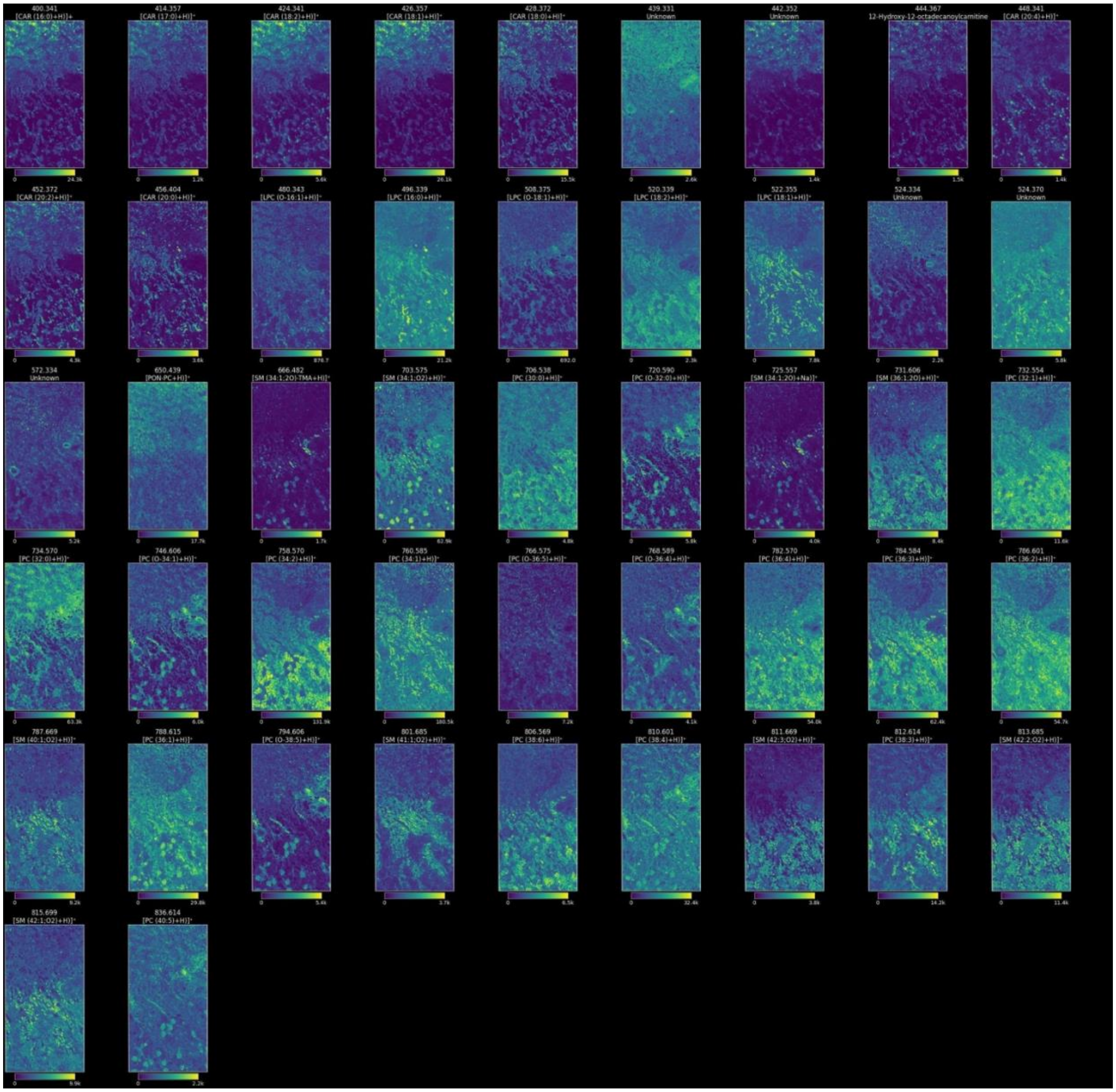

***Supplementary Figure 8:*** *Mosaic of positive ion mode MALDI imaging mass spectrometry data generated from donor VAN0031.*

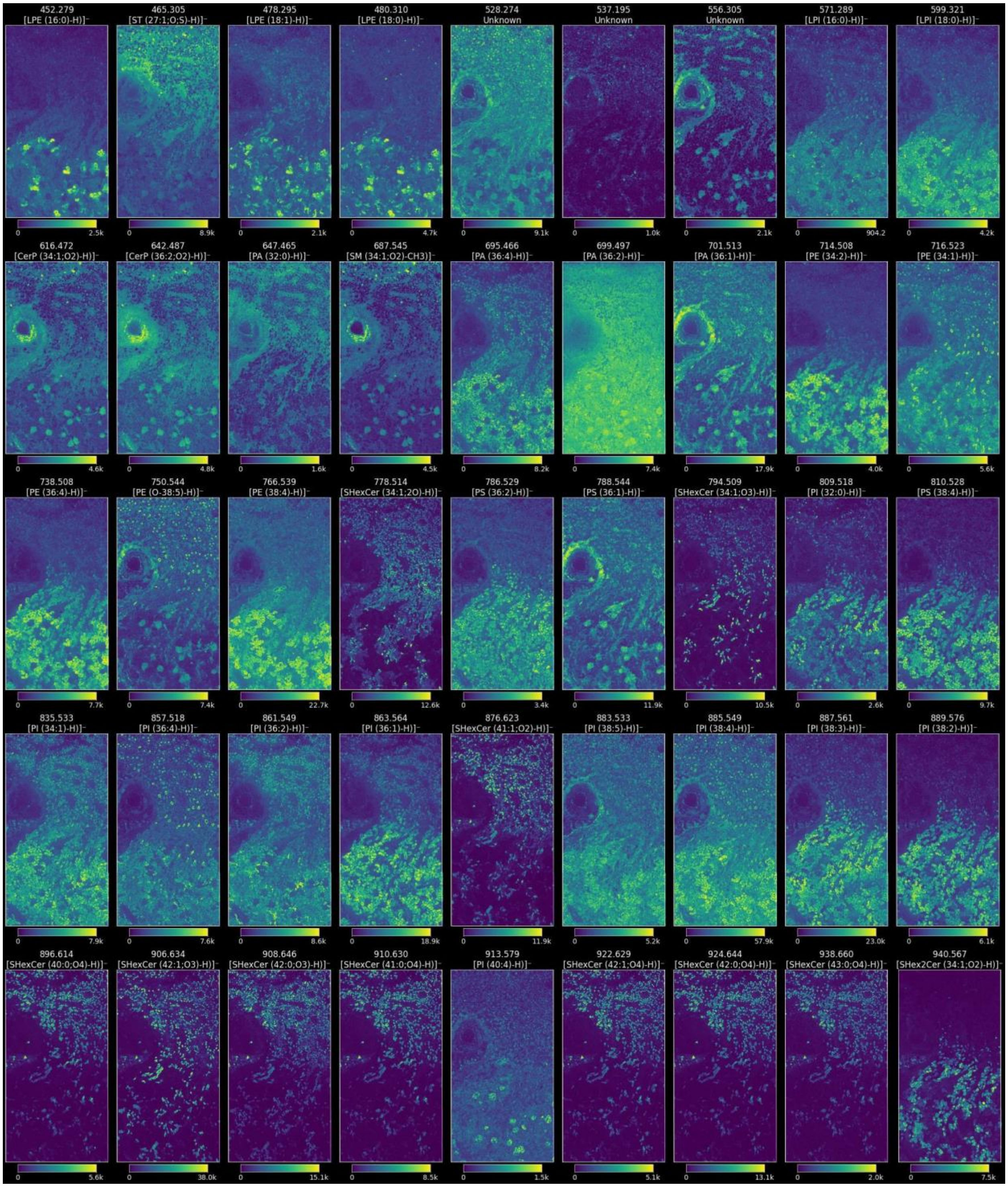


***Supplementary Figure 9:*** *Mosaic of negative ion mode MALDI imaging mass spectrometry data generated from donor VAN0031.*

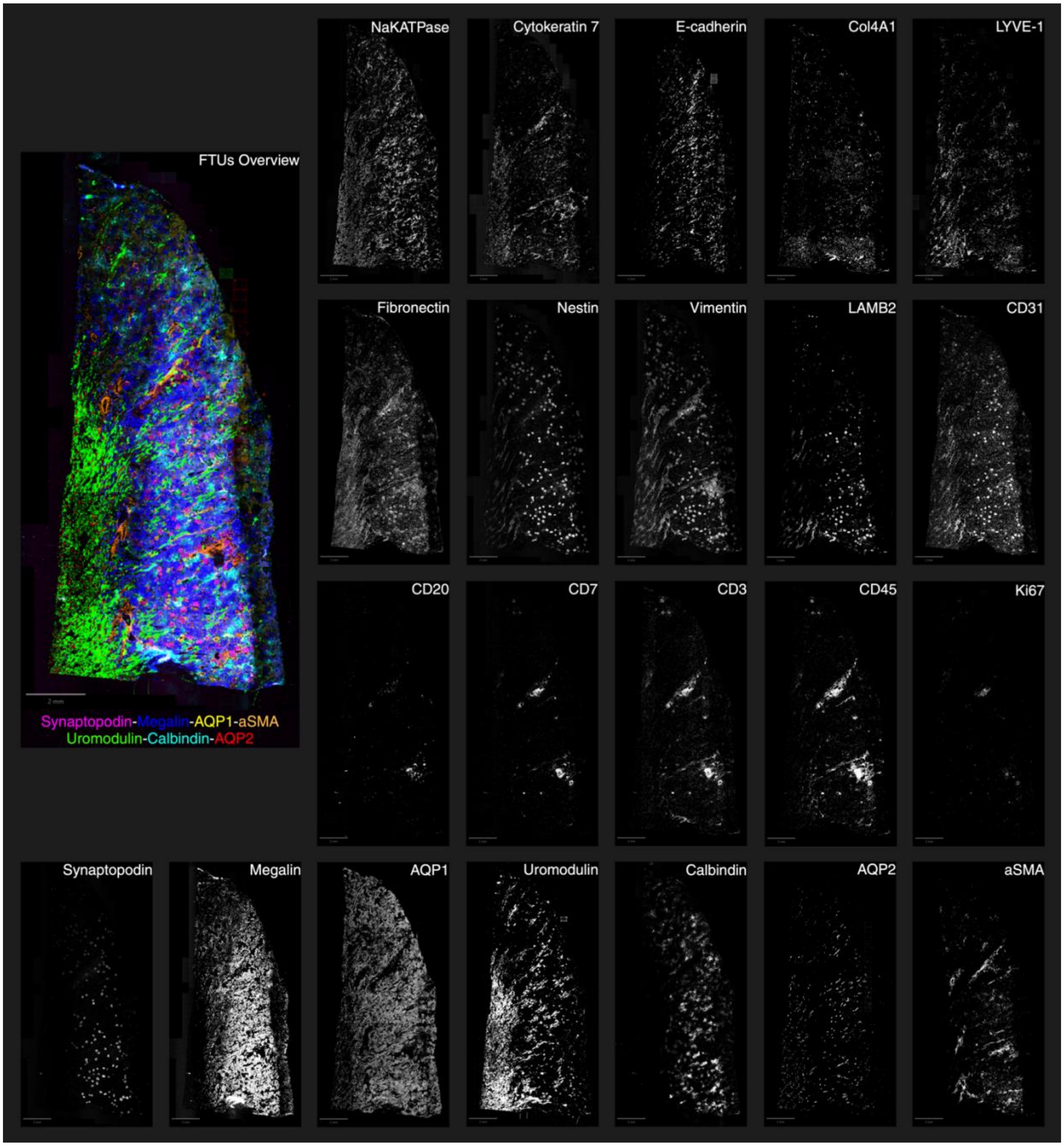


***Supplementary Figure 10:*** *CODEX multiplexed immunofluorescence microscopy images generated from donor VAN0005. The CODEX antibody panel includes 22 markers. An overview of Functional Tissue Units (FTUs) images is included with multiple markers overlaid in distinct colors: Synaptopodin (purple), Megalin (blue), AQP1 (yellow), aSMA (orange), Uromodulin (green), Calbindin (cyan), and AQP2 (red) (top left). Individual antibody staining are shown in grayscale for each marker. Scale bar: 2 mm.*

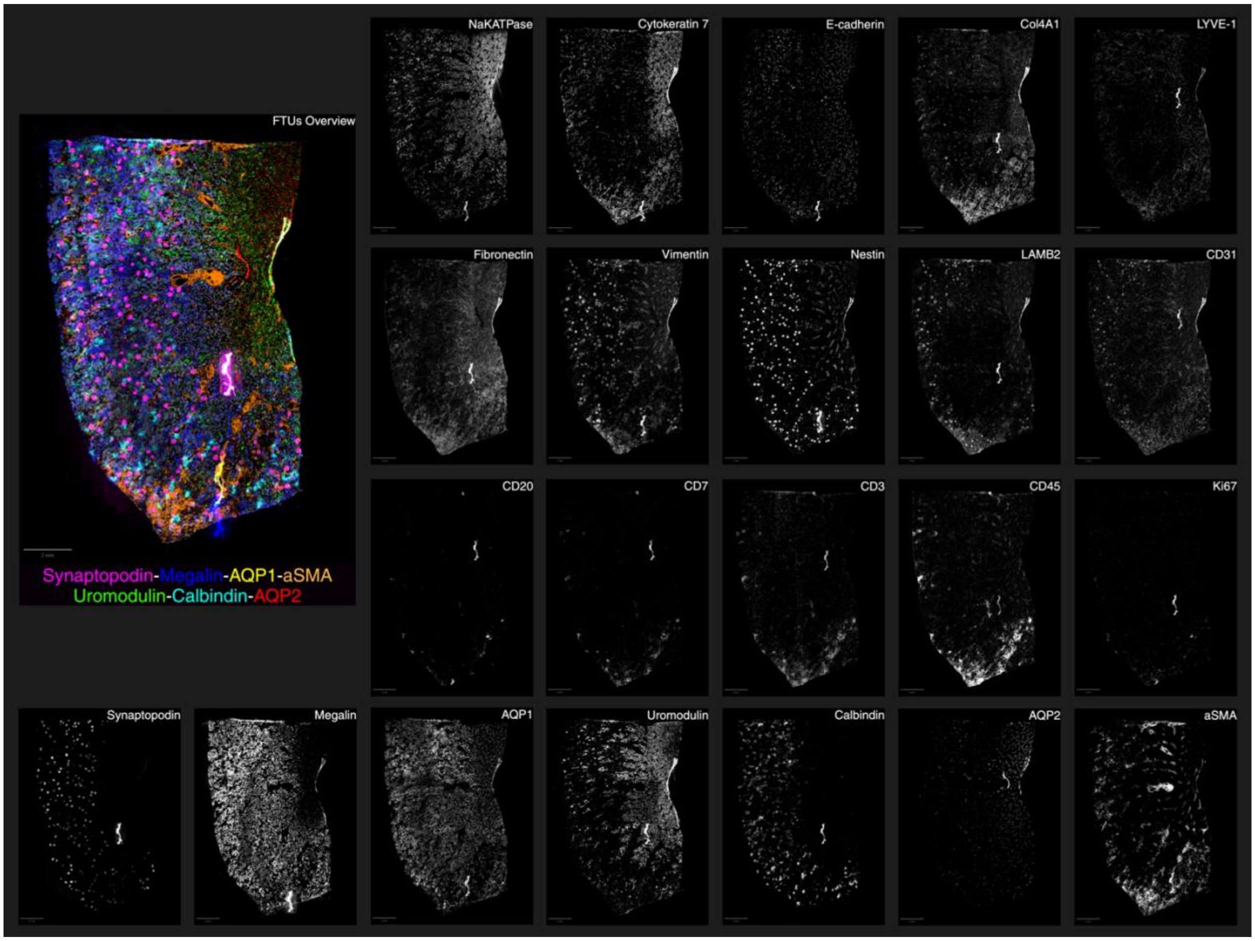


***Supplementary Figure 11:*** *CODEX multiplexed immunofluorescence microscopy images generated from donor VAN0019. The CODEX antibody panel includes 22 markers. An overview of Functional Tissue Units (FTUs) images is included with multiple markers overlaid in distinct colors: Synaptopodin (purple), Megalin (blue), AQP1 (yellow), aSMA (orange), Uromodulin (green), Calbindin (cyan), and AQP2 (red) (top left). Individual antibody staining are shown in grayscale for each marker. Scale bar: 2 mm.*

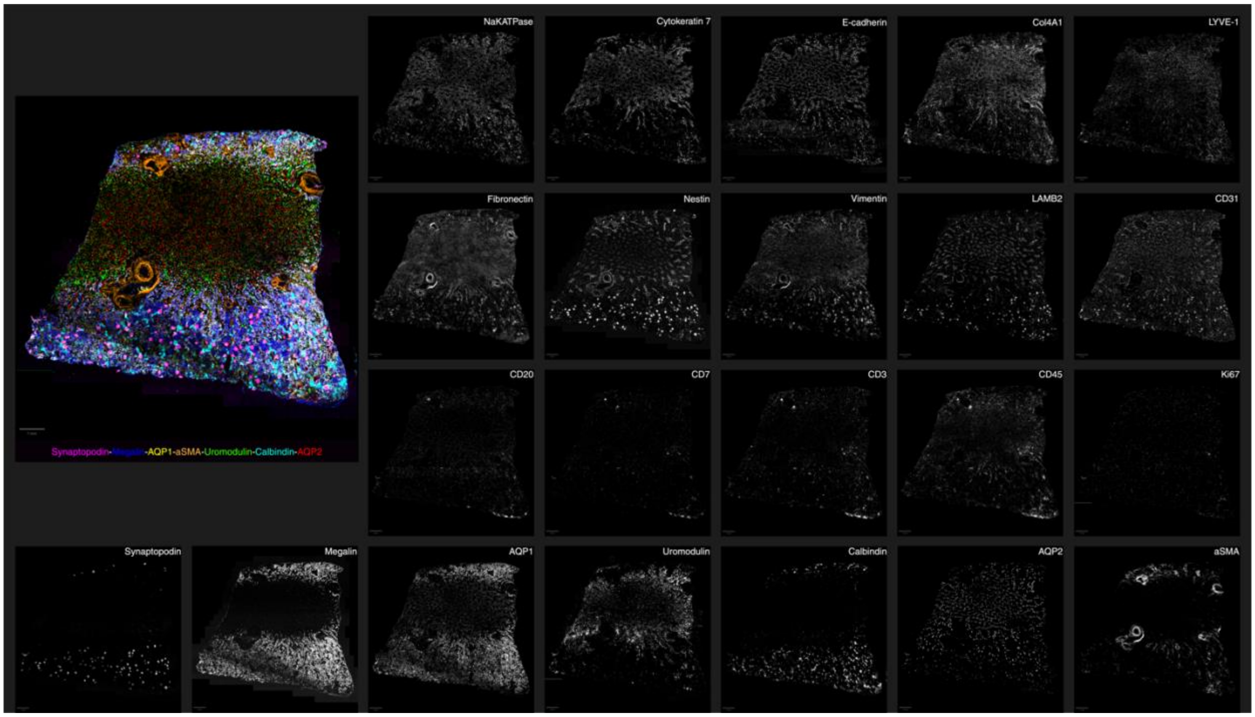


***Supplementary Figure 12:*** *CODEX multiplexed immunofluorescence microscopy images generated from donor VAN0031. The CODEX antibody panel includes 22 markers. An overview of Functional Tissue Units (FTUs) images is included with multiple markers overlaid in distinct colors: Synaptopodin (purple), Megalin (blue), AQP1 (yellow), aSMA (orange), Uromodulin (green), Calbindin (cyan), and AQP2 (red) (top left). Individual antibody staining are shown in grayscale for each marker. Scale bar: 1 mm.*

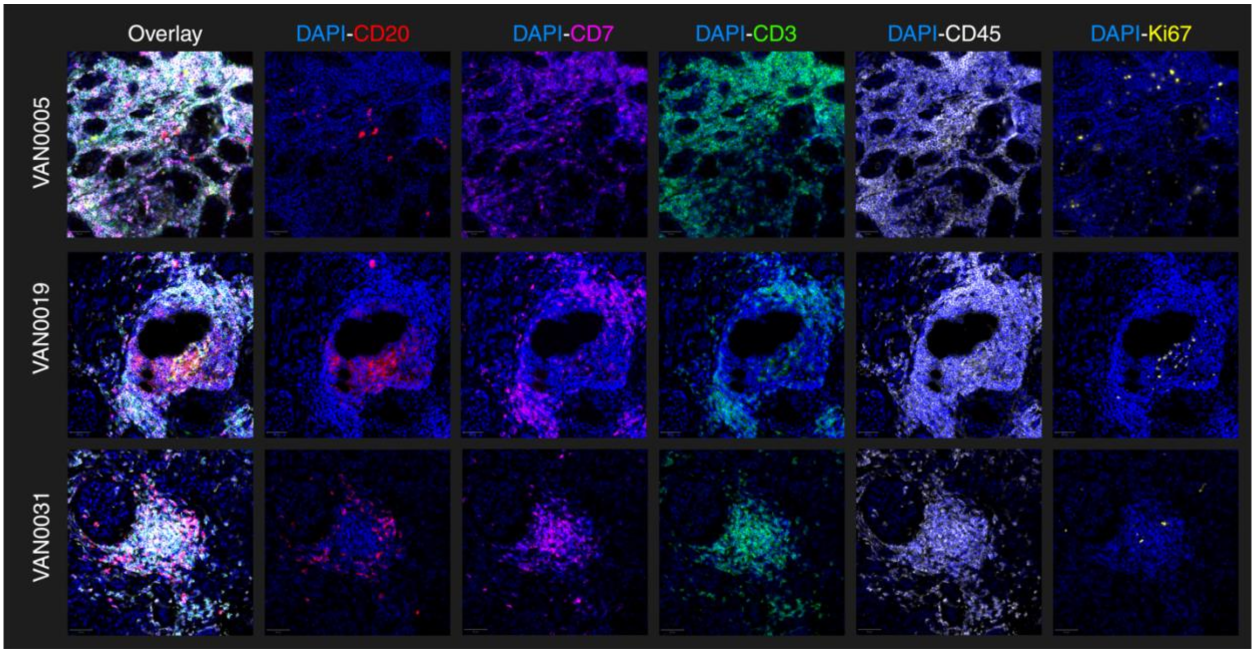


***Supplementary Figure 13:*** *Zoomed in CODEX multiplexed immunofluorescence microscopy images from a region of immune cell infiltrate. Selected immune markers are overlaid with the DAPI channel from kidney tissue samples from three donors (VAN0005, VAN0019 and VAN0031). Scale bar: 50 µm*

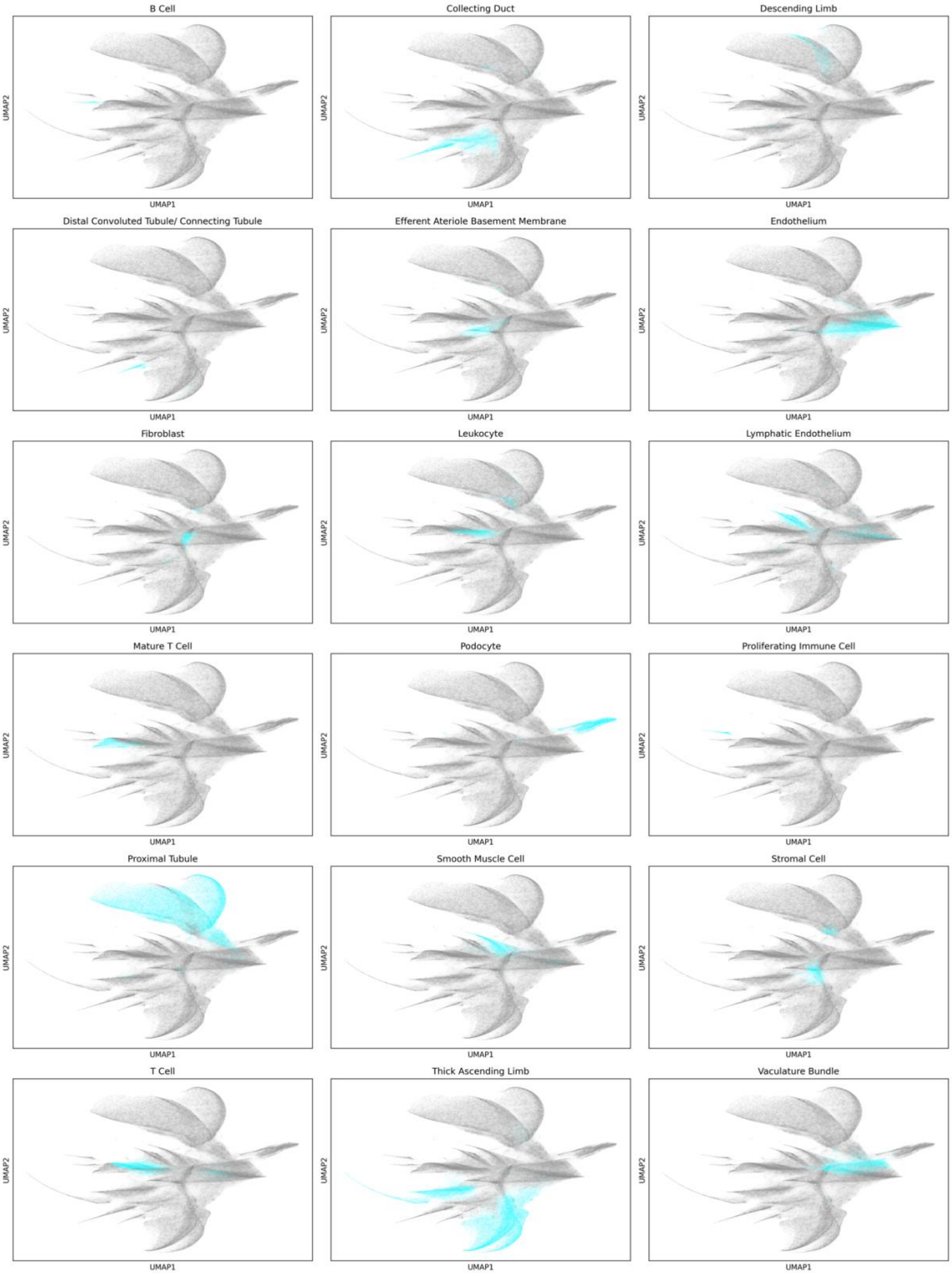


***Supplementary Figure 14:*** *Grid of UMAP plots displaying individual cell types, with each type highlighted in blue and other cells shown in gray. Each subplot represents a distinct cell type, illustrating its distribution in UMAP space.*

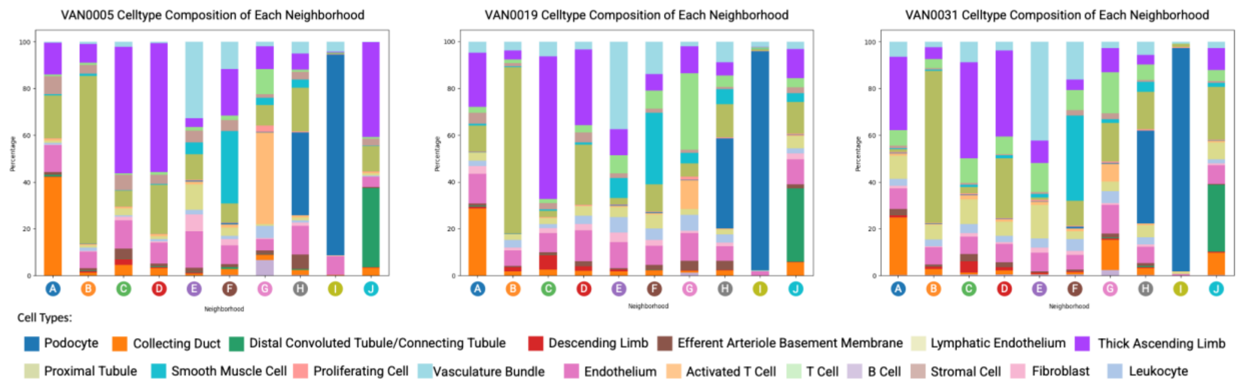


***Supplementary Figure 15:*** *Bar plots showing the cell type composition of each neighborhood in samples VAN0005, VAN0019, and VAN0031. Each bar represents a neighborhood, with different colors indicating the proportional abundance of distinct cell types.*

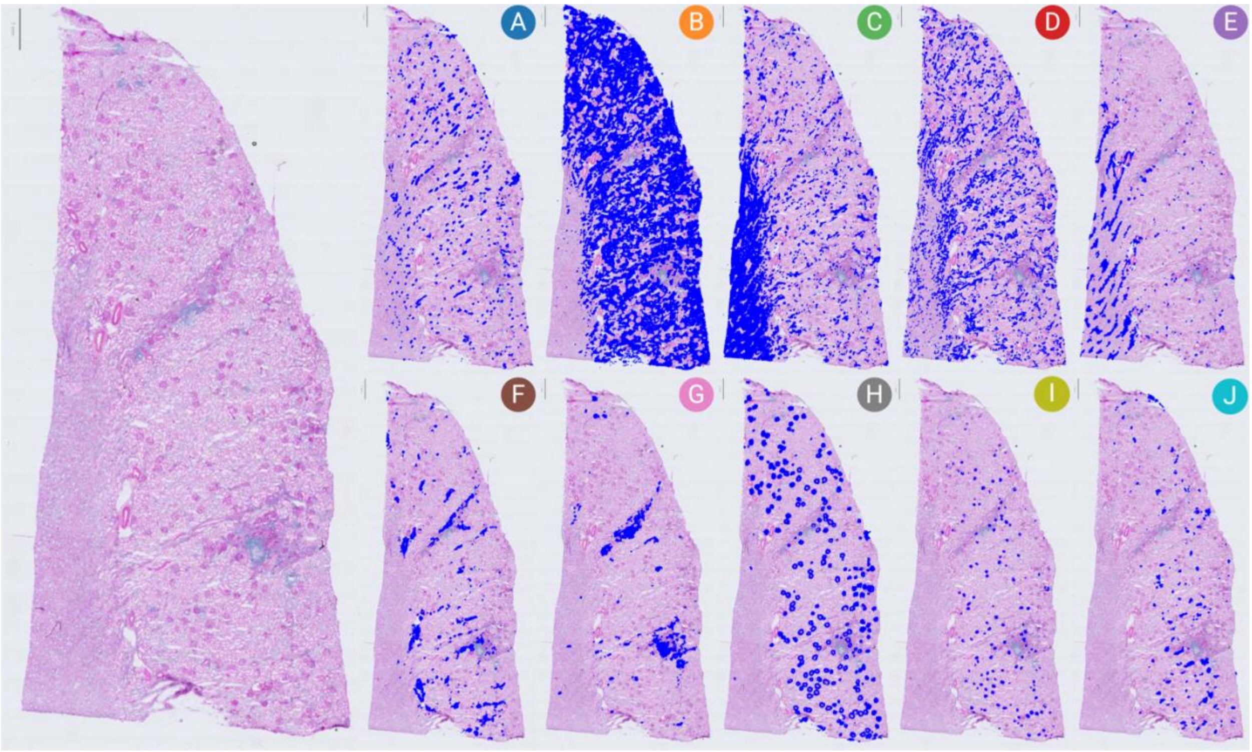


***Supplementary Figure 16:*** *Periodic acid-Schiff (PAS) stained image of human kidney tissue from donor VAN0005. CODEX-derived cellular neighborhoods are shown in a blue overlay on the PAS-stained image (Left). Each panel (A-J) represents different neighborhoods. Scale bar: 1mm.*

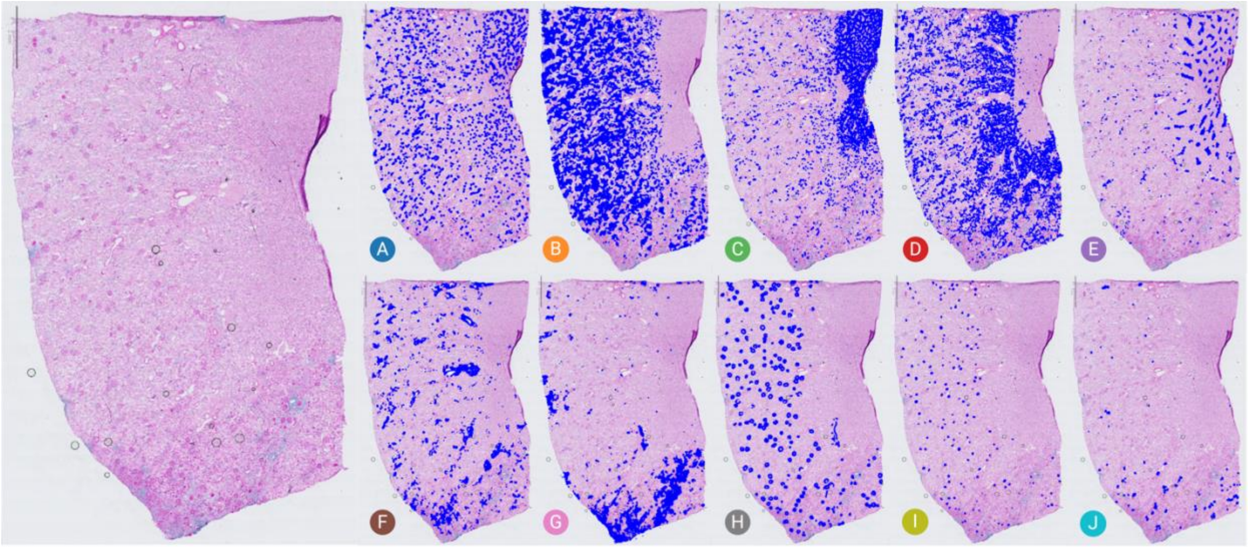


***Supplementary Figure 17:*** *Periodic acid-Schiff (PAS) stained image of human kidney tissue from donor VAN0019. CODEX-derived cellular neighborhoods are shown as a blue overlay on the PAS-stained image (Left). Each panel (A-J) represents different neighborhoods. Scale bar: 2mm.*

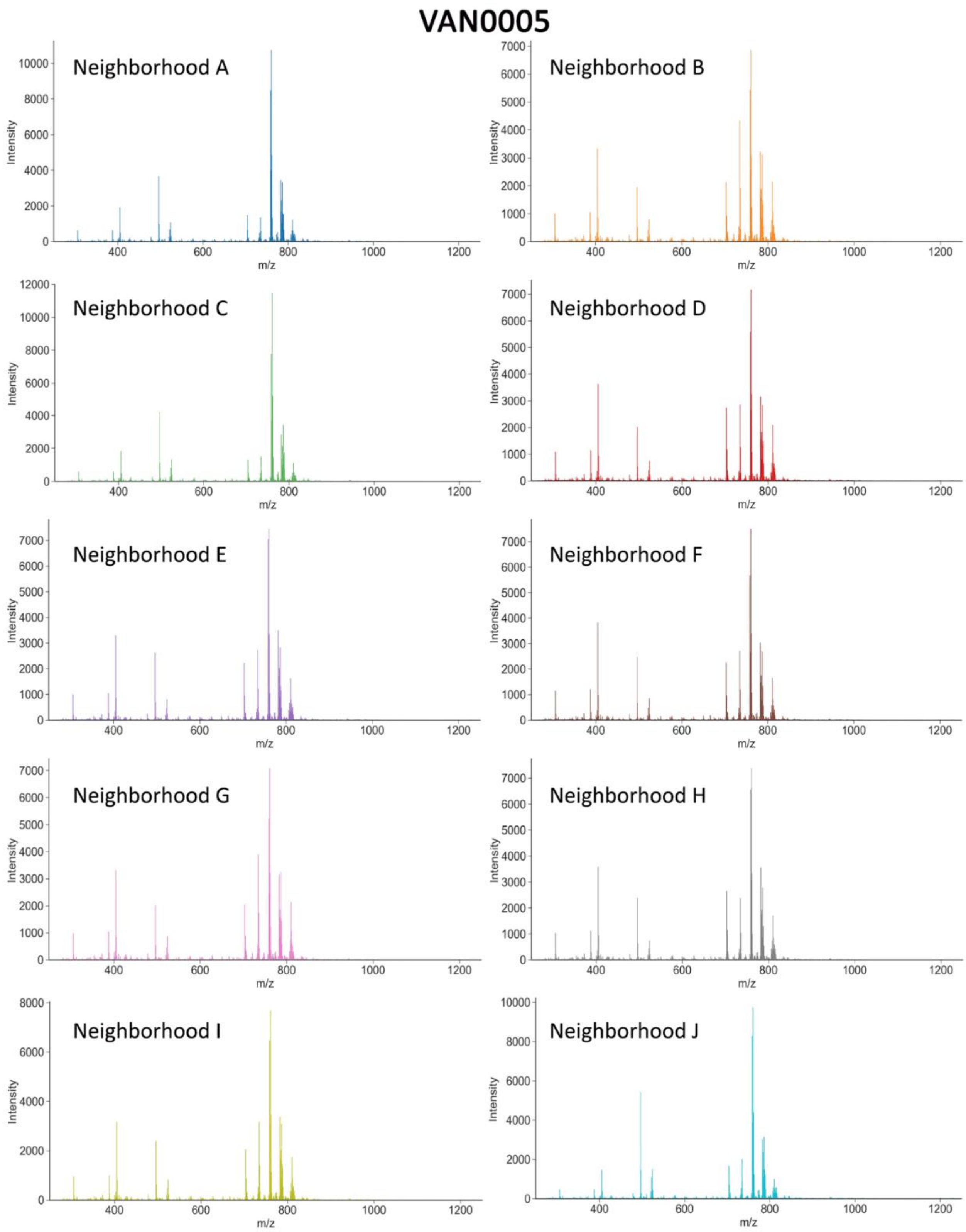


***Supplementary Figure 18:*** *Average mass spectrum of positive ion mode MALDI imaging mass spectrometry data generated from donor VAN0005.*

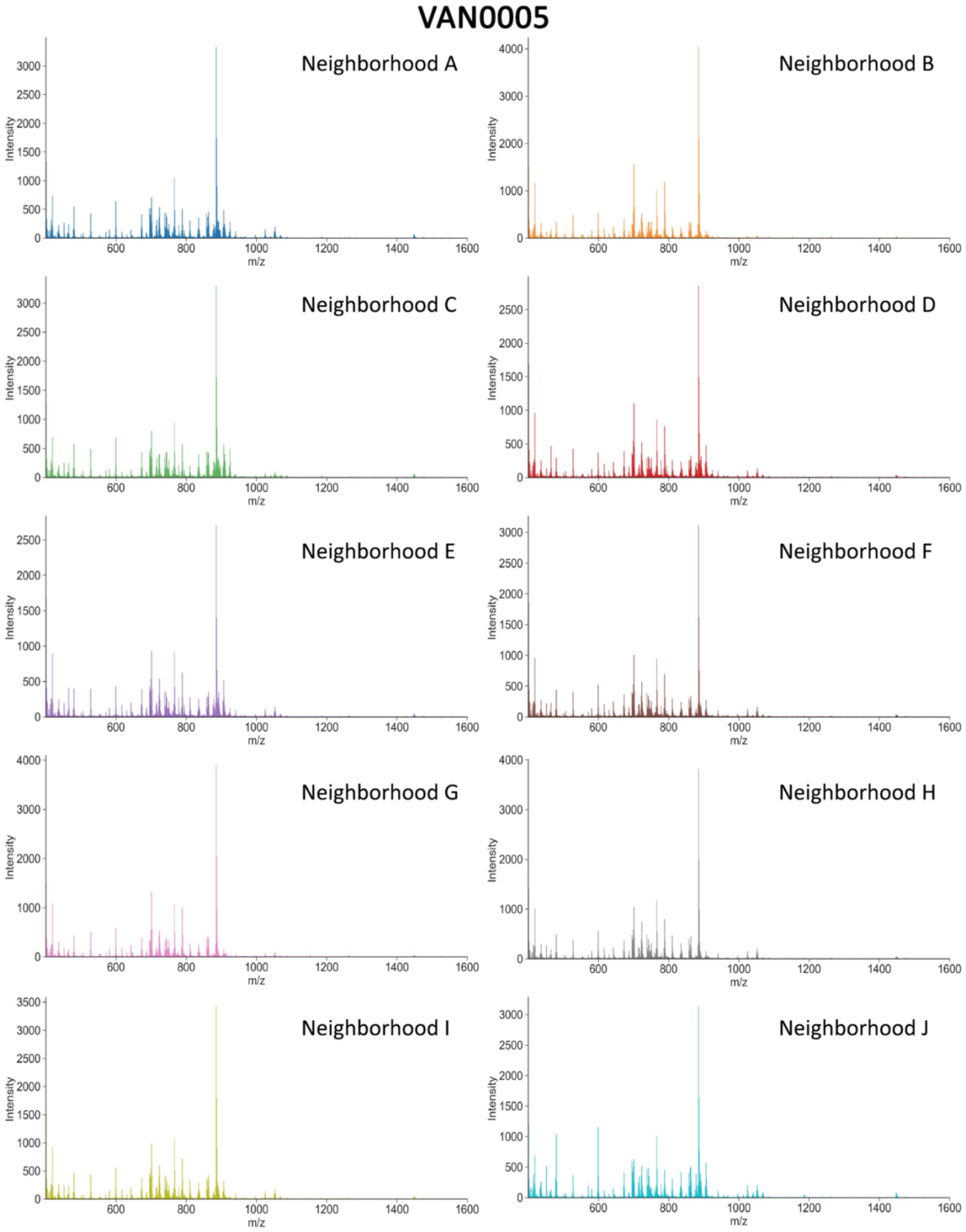


***Supplementary Figure 109:*** *Average mass spectrum of negative ion mode MALDI imaging mass spectrometry data generated from donor VAN0005*

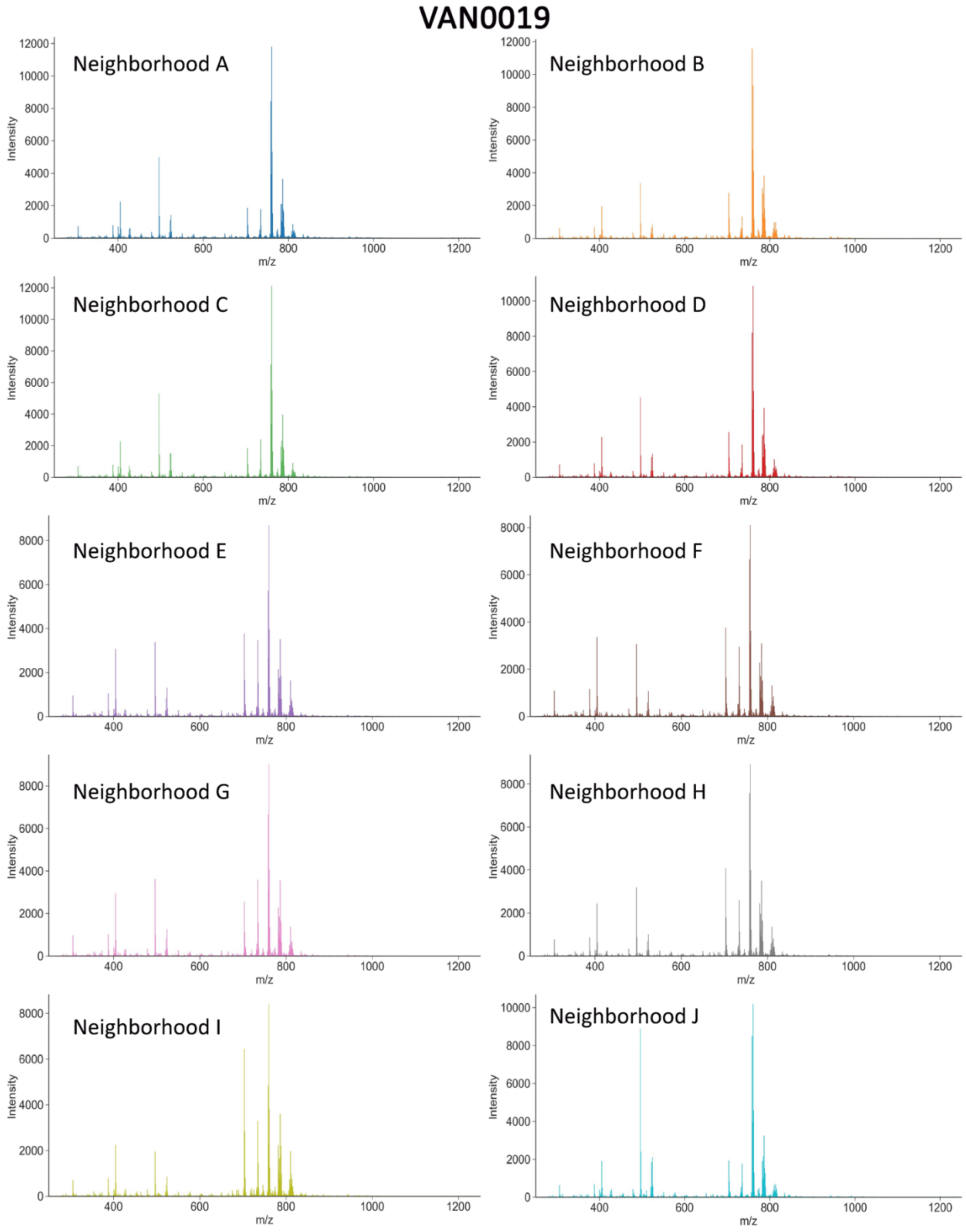


***Supplementary Figure 20:*** *Average mass spectrum of positive ion mode MALDI imaging mass spectrometry data generated from donor VAN0019.*

VAN0019

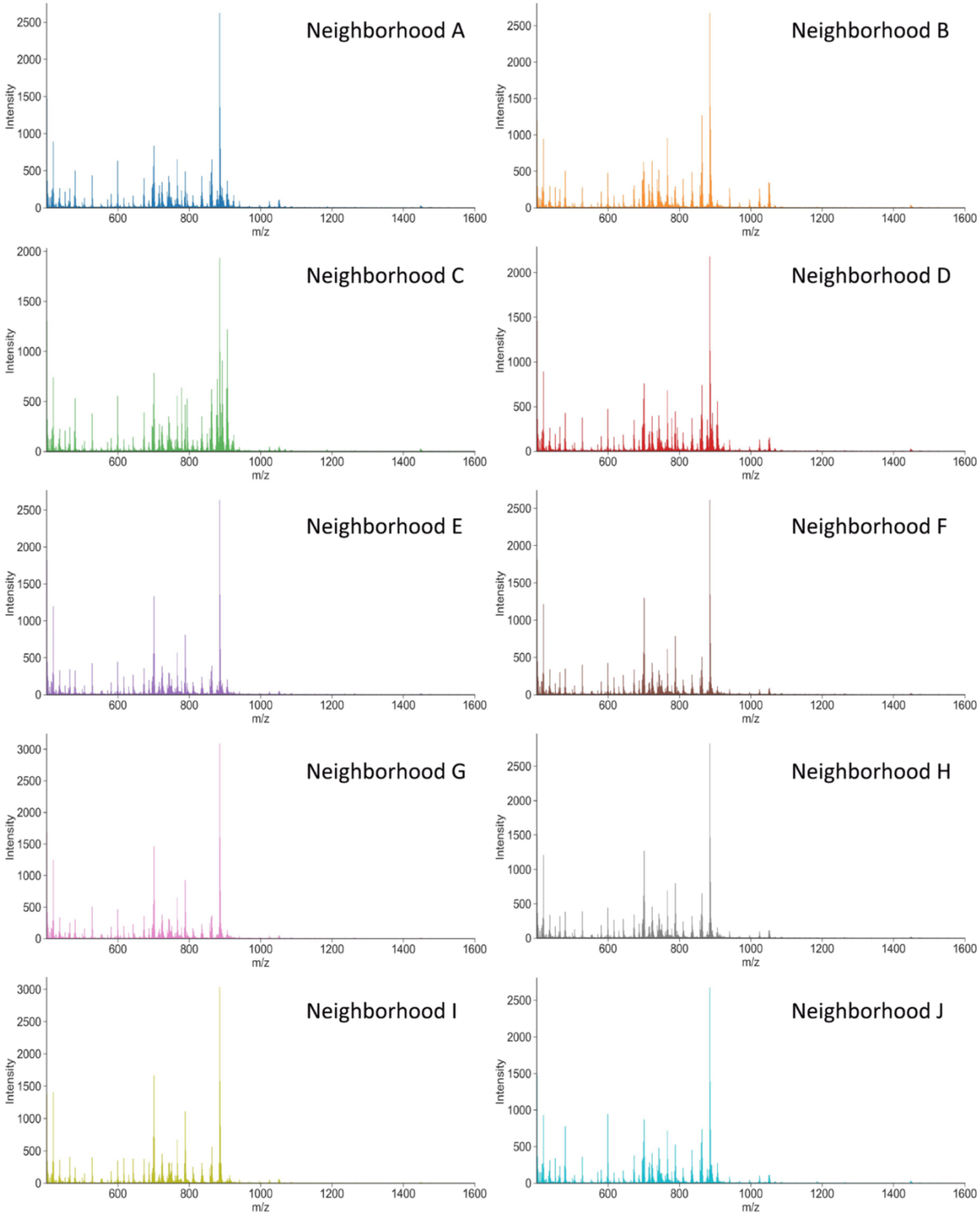


***Supplementary Figure 21:*** *Average mass spectrum of negative ion mode MALDI imaging mass spectrometry data generated from donor VAN00019*

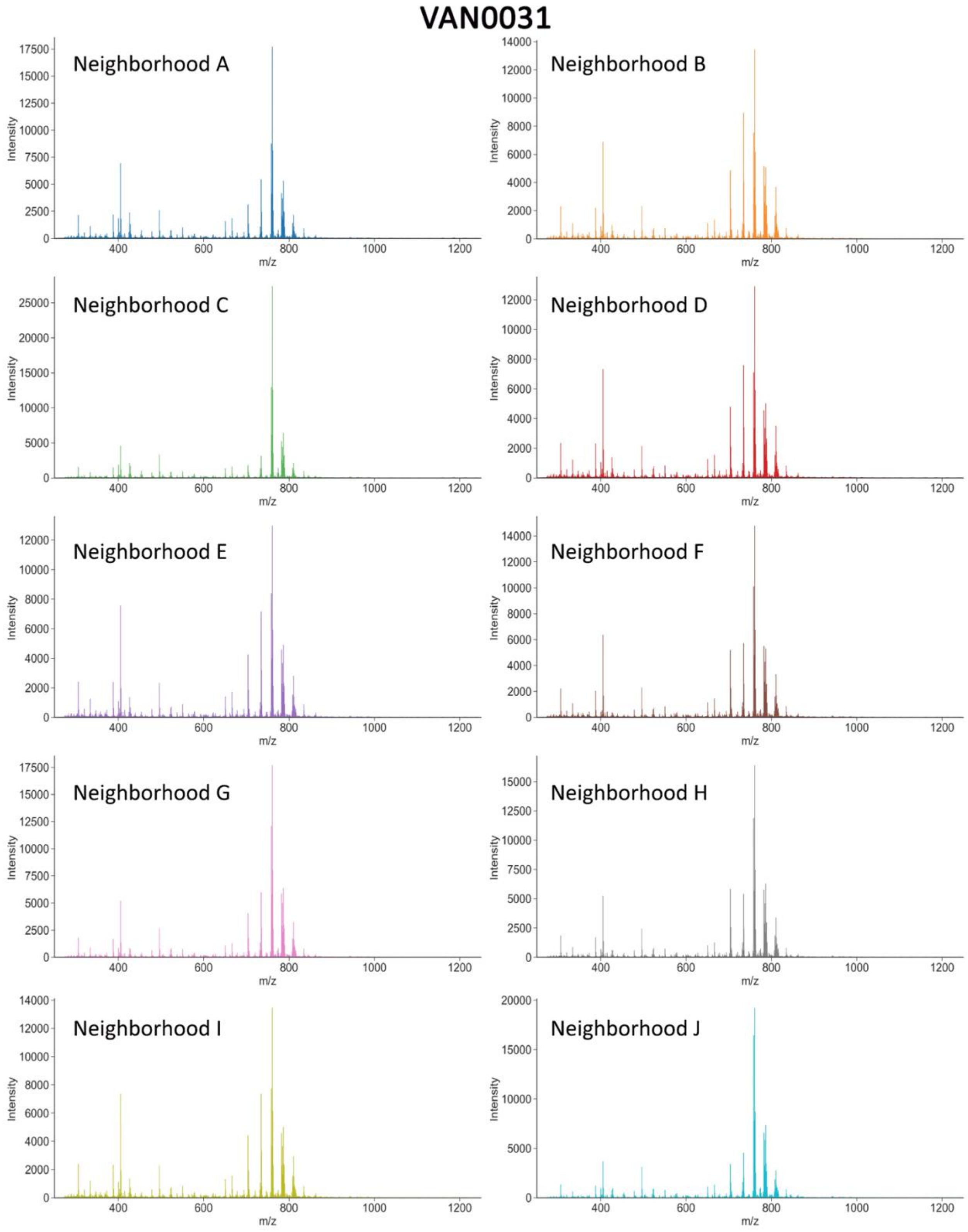


***Supplementary Figure 22:*** *Average mass spectrum of positive ion mode MALDI imaging mass spectrometry data generated from donor VAN0031*.

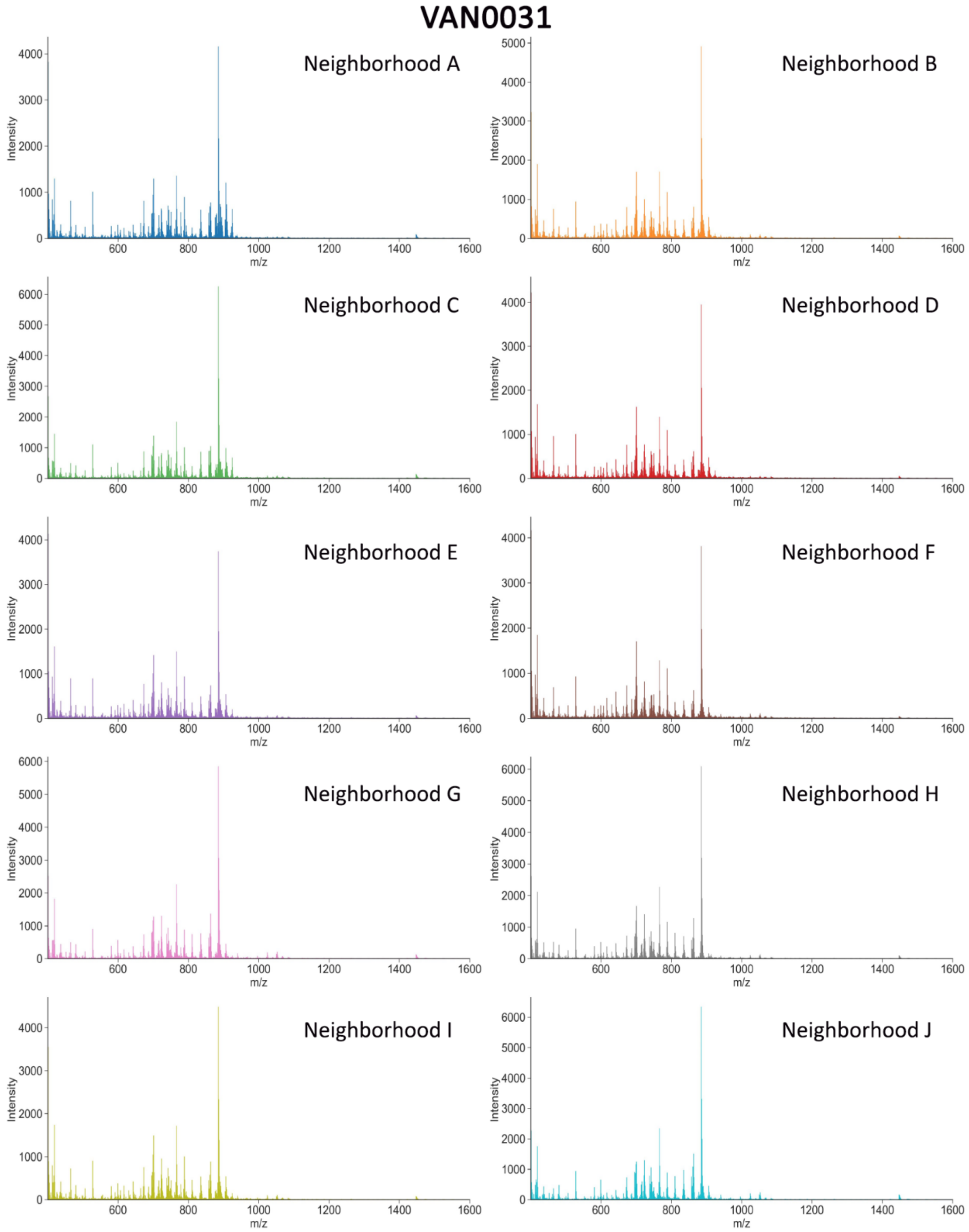


***Supplementary Figure 23:*** *Average mass spectrum of negative ion mode MALDI imaging mass spectrometry data generated from donor VAN0031*

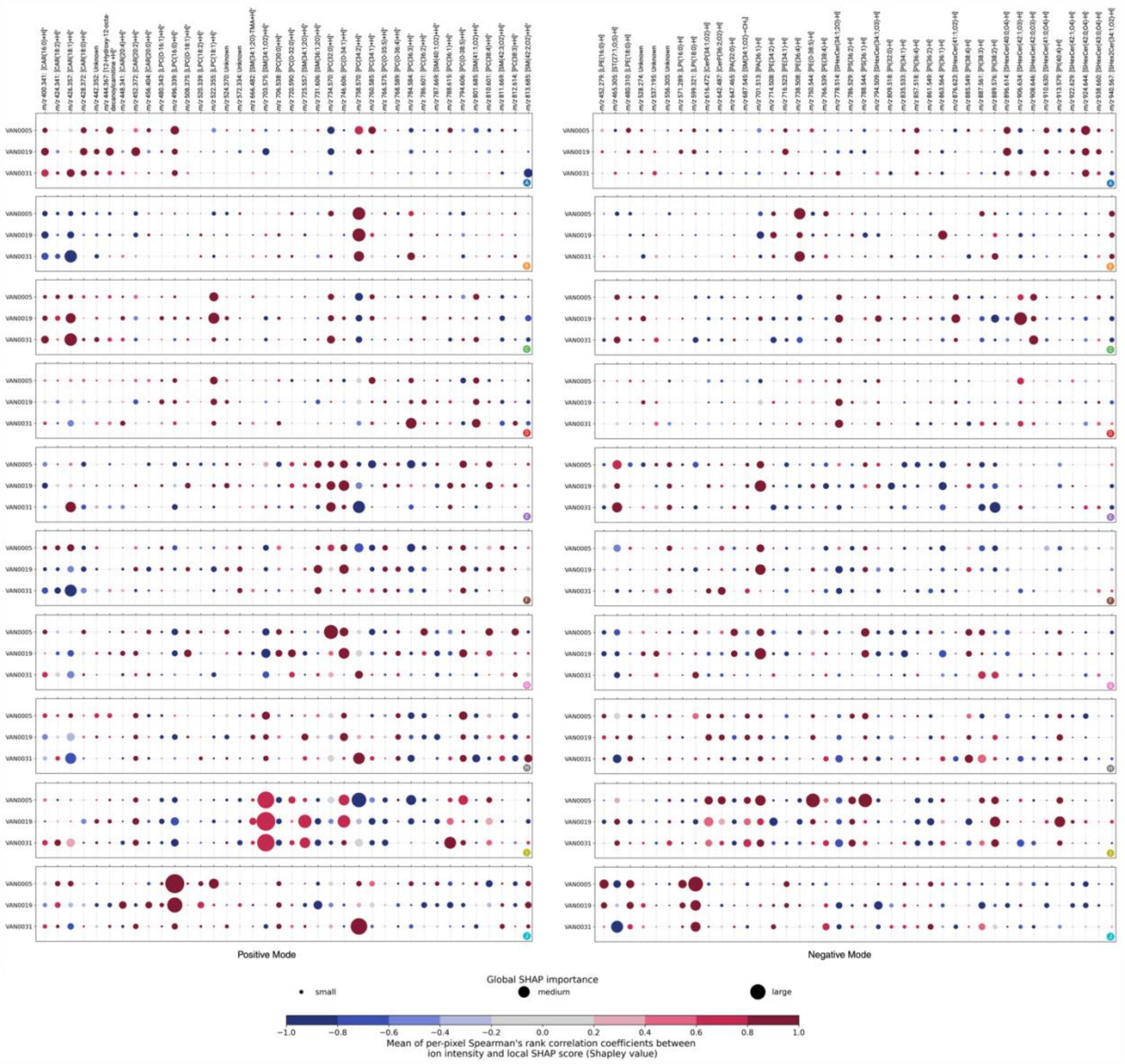


***Supplementary Figure 11:*** *Summary of neighborhood-specific biomarker candidates in positive (left) and negative (right) ion modes, obtained by applying our SHAP-based workflow to IMS-CODEX data. The columns correspond to a selection of molecular species (in increasing order of mass-to-charge ratios) that are biomarker candidates for each neighborhood. The rows correspond to different donors. Each bubble marker is informative of the direction (positive or negative correlation) and magnitude (relatively large or small) of a molecular species' influence on the classification model designed to recognize one of the 10 CODEX-derived neighborhoods (A-J). The marker size represents the magnitude of the molecular species' influence, as measured by its tissue sample-wide SHAP importance score for a given donor sample. The marker color indicates the direction of the molecular species' influence, as measured by the Spearman's rank correlation coefficient between the molecular species' mean-centered ion intensity values and its local pixel-specific SHAP scores. A positive Spearman's rank correlation coefficient indicates that a high ion intensity of the molecular species correlates with the neighborhood. Conversely, a negative Spearman's rank correlation coefficient indicates that a low ion intensity of the molecular species correlates with the neighborhood.*
